# Paf1C denoises transcription and growth patterns to achieve organ shape reproducibility

**DOI:** 10.1101/2022.03.25.485770

**Authors:** Duy-Chi Trinh, Marjolaine Martin, Lotte Bald, Alexis Maizel, Christophe Trehin, Olivier Hamant

## Abstract

In multicellular systems, all cells exhibit transcriptional noise. However, its exact contribution to morphogenesis often remains unclear, especially in animals where cells can also move. Here we take advantage of walled plant cells, where transcriptional noise happens in tissues with a fixed topology. Using synchronously growing guard cells in stomata, we demonstrate that mutation in *VIP3*, a subunit of the conserved polymerase-associated factor 1 complex (Paf1C), increases transcriptional noise in Arabidopsis. This conclusion could be generalized to other group of cells at the shoot apex. Such noise translates into growth and shape defects. Indeed, in *vip3* sepals, we measured higher growth heterogeneity between adjacent cells, with molecular evidence of increased local mechanical conflicts. This even culminated with the presence of negatively growing cells in specific growth conditions. Interestingly, such increased local noise makes the regional pattern of growth less clear-cut. Reduced regional conflicts lead to delay in organ growth arrest, ultimately making final organ shapes and sizes more variable. We propose that transcriptional noise is managed by Paf1C to ensure organ robustness by building up mechanical conflicts at the regional scale, instead of the local scale.

**HIGHLIGHTS:**

- Paf1C controls transcriptional noise in Arabidopsis
- Paf1C reduces growth heterogeneity and dampens local mechanical conflicts
- Impairing both Paf1C and microtubules can trigger negative growth in Arabidopsis
- Scaling up local conflicts to regional ones with sharp boundaries contributes to organ shape robustness

## INTRODUCTION

Stochastic gene expression is increasingly reported in living organisms, from bacteria growing in the same Petri dish (Elowitz et al., 2002) to multicellular organisms, such as Drosophila (Little et al., 2013) or Arabidopsis (Araújo et al., 2017; Meyer et al., 2017). The instructive role of such noise is increasingly studied in cell biology, notably in relation to developmental robustness (Nicholson, 2019; Tsimring, 2014). For instance, transcriptional noise can be exploited by cells for fate exploration and commitment (Chang et al., 2008; Desai et al., 2021; Weinberger et al., 2005). Notch-dependent cell fate in Drosophila is progressively acquired over time, largely through initially stochastic processes (Heitzler & Simpson, 1991). Similar fate acquisition from initially fluctuating expression levels has been reported in Arabidopsis (Meyer et al., 2017). Weak signals can also become unmasked thanks to stochastic resonance, *i.e.* adding noise makes them pass the threshold of detection (Rué et al., 2012). At a more integrated level in both space and time, fluctuation in cellular growth rate and orientation allows its averaging over time, and this contributes to organ shape reproducibility in Arabidopsis (Hong et al., 2016).

Whereas noise can contribute to patterning and morphogenesis in a bottom-up way, it is also classically thought to be channeled by global cues, notably to ensure shape robustness. For instance, precise timing of organ initiation reduces organ shape variability in Arabidopsis (Zhu et al., 2020). Morphogen gradients synchronize heterogeneous cell populations, and can even coordinate growth arrest through dilution as the organ becomes larger (Boulan & Léopold, 2021). Mechanical stress fields have been proposed to provide additional synchronizing cues (Aegerter-Wilmsen et al., 2007; Hufnagel et al., 2007; Shraiman, 2005). In particular, differential growth between domains has been proposed to trigger mechanical conflicts, ultimately leading to growth arrest in Drosophila (Pan et al., 2016) and Arabidopsis (Hervieux et al., 2016).

Beyond growth and development, noise management ultimately applies to transcriptional control. The causes of noise in gene expression are numerous, from chromatin modifications (Raser & O’Shea, 2004) to DNA topology (Elowitz et al., 2002) or ribosomal activity (Guido et al., 2007). More fundamentally, transcriptional noise is a product of the gene network topology (Acar et al., 2005), also consistent with the idea that “the cell is not a machine” (Nicholson, 2019). Interestingly, a genetic screen considering transcriptional noise as a quantitative trait identified the progression of transcriptional elongation phase by RNA polymerase II as a leading cause of noise in gene expression in yeast. This notably involves the polymerase-associated factor 1 complex (Paf1C) (Ansel et al., 2008). Whether this also applies to multicellular systems is unknown and is the focus of the present study.

In Eukaryotes, Paf1C is recruited to active Pol II elongation machinery. Through different mechanisms, Paf1C regulates multiple aspects of transcription including elongation and termination, as well as histone modification and chromatin structure (Tomson & Arndt, 2013; Van Oss et al., 2017). The roles of Paf1C are systemic. For example, the complex regulates transcriptional elongation of nearly all genes in yeast (Fischl et al., 2017; Hou et al., 2019). Yeast and mammalian Paf1Cs consist of five and six subunits respectively, and their six homologs in Arabidopsis have been identified (Antosz et al., 2017; Oh et al., 2004; Zhang et al., 2003) (Figure S1A). Mutants in the Arabidopsis Paf1C subunits were first uncovered thanks to their early flowering phenotype (Zhang et al., 2003). Paf1C mutants exhibit other developmental defects such as insensitivity to touch, increased irregularity in phyllotaxis and indeterminacy in flower development (Fal et al., 2017, 2019; Jensen et al., 2016) (Figure S1B). Despite this extensive biochemical knowledge and phenotypic characterization on Paf1C in multicellular systems, whether transcriptional noise is involved and whether noise could relate to these phenotypic features are unknown.

To link transcriptional noise and developmental robustness, we used a mutant of *VIP3* which encodes for a component of the Paf1C in Arabidopsis, with a focus on sepals, the outermost floral organs. Arabidopsis sepals are highly robust in shape and size, making them an ideal model system to investigate the nexus between the intrinsic control of shape robustness and transcriptional noise (Roeder, 2021). Using several reporter lines in tissues with stereotyped and iterative developments (stomata and shoot meristems), we confirmed that transcriptional noise is indeed increased in the Paf1C mutant. Long term imaging of sepal growth also revealed that growth in the mutant is more variable locally. We find that increased local growth heterogeneity hinders the formation of global consistent domains with distinct growth properties. This diffuses mechanical conflicts to a more local scale, between adjacent cells instead of between regions. In the end, sepals fail to arrest their growth, subsequently leading to more variable final organ shapes and sizes.

## RESULTS

### Mutation in *VIP3* increases the variability in linker histone H1 expression between sister guard cells

Stomata are epidermal valves formed by a pair of specialized guard cells. These guard cells originate from a common guard mother cell following a symmetric division (Figure 1A). Because two sister guard cells are small and close to each other, they likely experience very similar internal and external cues. Given their common developmental origin, close position and synchronous differentiation, guard cells offer an excellent system to study gene expression variability.

**Figure 1.**
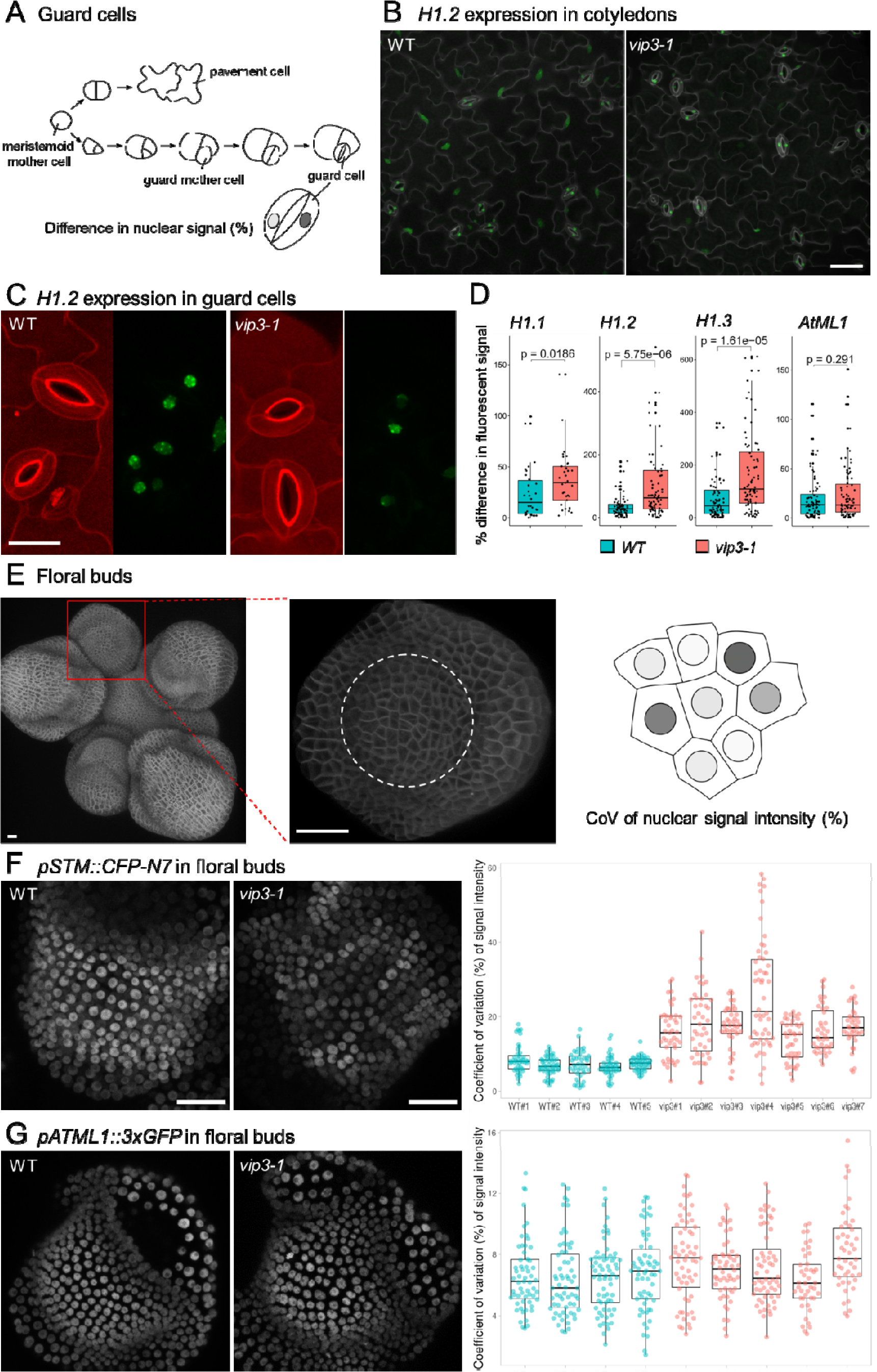
Gene expression variability in guard cells and floral buds in WT and *vip3-1* (A) Stomatal development in Arabidopsis. Sister guard cells are formed through a symmetrical cell division. Using a fluorescent protein targeted to the nucleus, the difference in gene expression (nuclear signal) between the two guard cells can be calculated. Adapted from (Torii, 2007). (B) A large area of the cotyledon in WT and *vip3-1* showing the expression of *pH1.2::H1.2-eGFP* in guard cells. (C) Close-up, representative confocal images of stomatal guard cells and expression of *pH1.2::H1.2-eGFP* in WT and *vip3-1*. (D) Difference (in percentage) in fluorescent signal between two sister guard cells. *H1.1 (pH1.1::H1.1-eGFP), H1.2 (pH1.2::H1.2-eGFP), H1.3 (pH1.3::H1.3- eGFP), AtML1 (pAtML1::H2B-3xGFP::tRBCS)*. (E) A typical Arabidopsis inflorescence with the shoot apical meristem (SAM) in the center and several flower buds of different stages. Young flower buds are used to measure local variability in gene expression. Only cells in the center of the floral bud (white circle) are taken into account. Local variability in gene expression is measured as coefficient of variation (CoV% = SD/mean*100%) in nuclear signal intensity of a group of cells consisting of a cell and all of its contacting neighbours. (F) Representative image of *pSTM::CFP- N7* expression in stage 3 floral buds (left) and coefficient of variation (%) of signal intensity between neighbouring cells (right) for the WT and *vip3-1* samples. (G) Similar to (F), for *pAtML1::H2B-3xGFP::tRBCS*. Scale bars = 20 µm.

To do so, we used reporter lines for genes that are known to be expressed in guard cells. More specifically, we chose to analyze the expression of the whole H1 linker histone gene family: *H1.1 (pH1.1::H1.1-eGFP), H1.2 (pH1.2::H1.2-eGFP)* and *H1.3 (pH1.3::H1.3-eGFP)*. In normal conditions H1 genes are expressed in stomata and are required for stomata functioning (Rutowicz et al., 2015) (Figure 1B). Note that *H1.1* and *H1.2* exhibit a rather steady expression profile, whereas *H1.3* expression depends on stress level, including mechanical stress (Fal et al., 2021). We reasoned that this family was small enough to get an overview of their expression variability between WT and mutant, while being functionally relevant for stomata. We also included one of the regulators of epidermal cell identity *AtML1 (pAtML1::H2B-3xGFP::tRBCS),* which is expressed in all epidermal cells.

The use of GFP with a nuclear localization signal for AtML1 or translational fusion for histone markers allowed us to compare expression levels using total signal in nuclei as a proxy. To assess gene expression variability of these reporter lines in a pair of guard cells, we acquired confocal images of stomata in seedling cotyledons, and calculate the difference in nuclear signal intensity (Δ, in %) between two sister guard cells, using the nucleus with weaker signal as reference (see Materials and methods). Similar trends were obtained when considering the average or the median, but here we only discuss the differences in median as the parameter is less sensitive to outliers.

Since the two *vip3* mutant alleles exhibit similar phenotypes (Figure S1C and S1D, (Fal et al., 2017, 2019; Jensen et al., 2016)), all the work in this study was performed on *vip3-1*. Figure 1C shows representative images of stomatal guard cells and *H1.2* expression in WT and *vip3-1*. Whereas the signal of *H1.1* and *H1.2* differed by 15% and 28% respectively in the WT, the figures for *vip3-1* were 34% and 63% respectively, *i.e.* double those of the WT (n_WT_(H1.1) = 30, n*_vip3-1_*(H1.1)=32, n_WT_(H1.2) = 80, n*_vip3-1_*(H1.2) = 80, *p*_H1.1_ = 0.019, *p*_H1.2_ = 5.75e^-06^, Kruskal–Wallis test) (Figure 1D). In normal growth condition, H1.3 was only expressed in a few guard cells in both backgrounds. We therefore placed 8-day old seedlings in dark for 24 hours to induce H1.3 expression as noted in (Rutowicz et al., 2015). This short period of darkness was sufficient to induce *H1.3* expression in cotyledon guard cells from both genotypes. In these conditions, *H1.3* expression in the WT appeared more variable between the pair of sister guard cells than H1.1 and H1.2 (⊗_WT_(H1.3) = 45%, *p_H1.1-H1.3_* = 0.0004, *p_H1.2- H1.3_* = 0.018, Kruskal–Wallis test). Furthermore, the variability of H1.3 expression in *vip3-1* was again roughly twice as high as in WT (Δ*_vip3-1_*(H1.3) =109%, n_WT_(H1.3) = 80, n*_vip3- 1_*(H1.3)=84, *p* = 1.61e^-05^, Kruskal–Wallis test) (Figure 1D). Note that we did not observe any significant difference in *AtML1* expression variability between sister guard cells in the WT and *vip3-1* (⊗_WT_(AtML1) = 13.4% *vs.* ⊗*_vip3-1_*(AtML1) = 13.1%, *p*= 0.291, Kruskal–Wallis test) (Figure 1D). This is consistent with the finding that in yeast Paf1C binds to RNA polymerase II at different degrees, depending on genes (Fischl et al., 2017). The rather stable *AtML1* expression level is also consistent with the finding that VIP3 is not affecting all pathways with the same strength in Arabidopsis (Fal et al., 2019), while also providing a baseline for our analysis. In short, using sister guard cells as stereotypical and synchronous cells, we show that H1 expression is locally more variable in *vip3-1*, when compared to the WT.

### Mutation in *VIP3* increases the local variability in gene expression between adjacent cells in floral and inflorescence meristems

Guard cells are an ideal system to analyze gene expression variability because they have comparable history. Yet, results may not be applicable to other tissues where adjacent cells undergo stochastic cell divisions. To challenge our finding, we thus assessed gene expression variability in other developmental contexts, including floral and inflorescence meristems (Figure 1E). Here we analyzed expression of *AGAMOUS (pAG::AG-2xVenus*), *SHOOT MERISTEMLESS* (*pSTM::CFP-N7*) and *ATML1 (pAtML1::H2B-3xGFP::tRBCS)*.

Because there are many cells in floral meristems and SAMs (as opposed to only a pair of guard cells), we calculated the coefficient of variation (CoV(%), defined as the standard deviation divided by the mean, in percentage) in gene expression within a group of adjacent cells (see Materials and methods) (Figure 1E, last diagram).

AG is a key regulator of reproductive organ development and is responsible for flower determinacy (Yanofsky et al., 1990). We previously showed that *AG* expression is likely to be affected in the *vip3-1* mutant and its mis-regulation has been correlated to the flower indeterminacy phenotype in the mutant (Fal et al., 2019). *AG* is expressed specifically in the center of a floral bud where inner organs (petals, stamens and carpels) will form. Here we worked with stage 3 floral buds where sepals have just emerged and not yet covered the buds (Smyth et al., 1990). Consistent with our previous report that *AG* signal was lower in *vip3-1* compared to the mutant (Fal et al., 2019), while most or all cells in the center in WT floral buds expressed *AG*, *vip3-1* floral buds usually lacked *AG* signal in some cells in the very center (Figure S1E, left). While the average CoV of local signal intensity of *AG* in the WT was 20.1%, the number for *vip3-1* was slightly, but significantly, higher, 28.6% (*p*=7.1e^-08^, Kruskal–Wallis test, n = 395 cells in 5 samples for WT, 352 cells in 7 samples for *vip3-1*) (Figure S1E, right).

*SHOOT MERISTEMLESS (STM)* regulates meristem functions and its expression defines meristematic tissues (Long et al., 1996). In floral buds of stage 2-3, *STM* expression is strong in the center but much weaker in emerging sepals. While nuclei in WT floral buds showed a homogenous signal, it was much less so in *vip3-1* with strong and weak nuclei beside each other (Figure 1F, left). We did a similar quantification of *pSTM::CFP-N7* signal intensity, and confirmed that *STM* signal varied just slightly between cells in the WT, with the average CoV of local signal intensity at 7.3% (5 samples, 224 cells). However, signal variability was more than double in the mutant, with the average CoV at 18.2% (7 samples, 295 cells, *p*=2.15e^-55^). Similar to the case of *AG*, the difference across samples in the mutant was more prominent than in the WT (Figure 1F, right).

*AtML1* expression has already been shown to be homogenous in floral meristems (CoV_WT_(AtML1) ∼ 10% according to (Meyer et al., 2017)). We examined *AtML1* expression levels in two tissues: inflorescence and stage 3 floral buds. In floral buds, while the local variation of *AtML1* intensity was just around 6.5%, the figure for *vip3-1* was slightly higher, 7.3% (*p*=0.00036, n_WT_ = 250 cells in 4 samples, n*_vip3-1_* = 266 cells in 5 samples) (Figure 1G). The situation is similar in SAMs, where local variation of *ATML1* intensity was slightly higher in the mutant compared to the WT (CoV_WT_ = 6.4% vs. CoV*_vip3-1_*=7.6%, *p*=3.3e^-13^, Kruskal–Wallis test, 3 samples each) (Figure S1F). Again, similar to the case of guard cells, *ATML1* gene expression variability in floral buds and SAMs is much lower than other genes studied.

In summary, working with various reporter lines in different tissue contexts (guard cells in cotyledons, group of adjacent cells in floral and inflorescence meristems), we demonstrated that mutation in *VIP3* leads to increased gene expression variability. Next, we investigated whether the increased local variability in gene expression in *vip3-1* may contribute to the global phenotype of the mutant.

### *VIP3* loss of function increases variability in abaxial sepal shape and size

*vip3* mutants have already been shown to exhibit variable phyllotaxis and floral termination (Figure S1A,B; (Fal et al., 2017, 2019). Because such complex phenotypes involve coordination between several organs, here we decided to focus on a simpler organ, the sepal. Each Arabidopsis flower produces four sepals, and each sepal exhibits a remarkably reproducible shape during its development. Thus, it is an ideal system to relate local noise and global shape robustness (Roeder, 2021).

We noticed that sepals in *vip3* mutants usually curve outwards and their shapes also appeared more variable than those in the WT (Figure 2A-B, Figure S1C). In theory, sepal shape variability may be an indirect consequence of patterning defects in flowers (Fal et al., 2019). To avoid such effects and to focus on the intrinsic control of sepal morphogenesis, we only selected flowers with a WT number of sepals (4 sepals) for all analyses. We also focused on abaxial sepals, which are the first to emerge from floral bud meristems in both genotypes (Figure 2C). To quantify the variability in *vip3-1* sepal shape, we employed the SepalContour tool described in (Hong et al., 2016). This tool allows us to extract several features of sepals, including sepal area, sepal contours and other shape descriptors such as sepal length and width.

**Figure 2.**
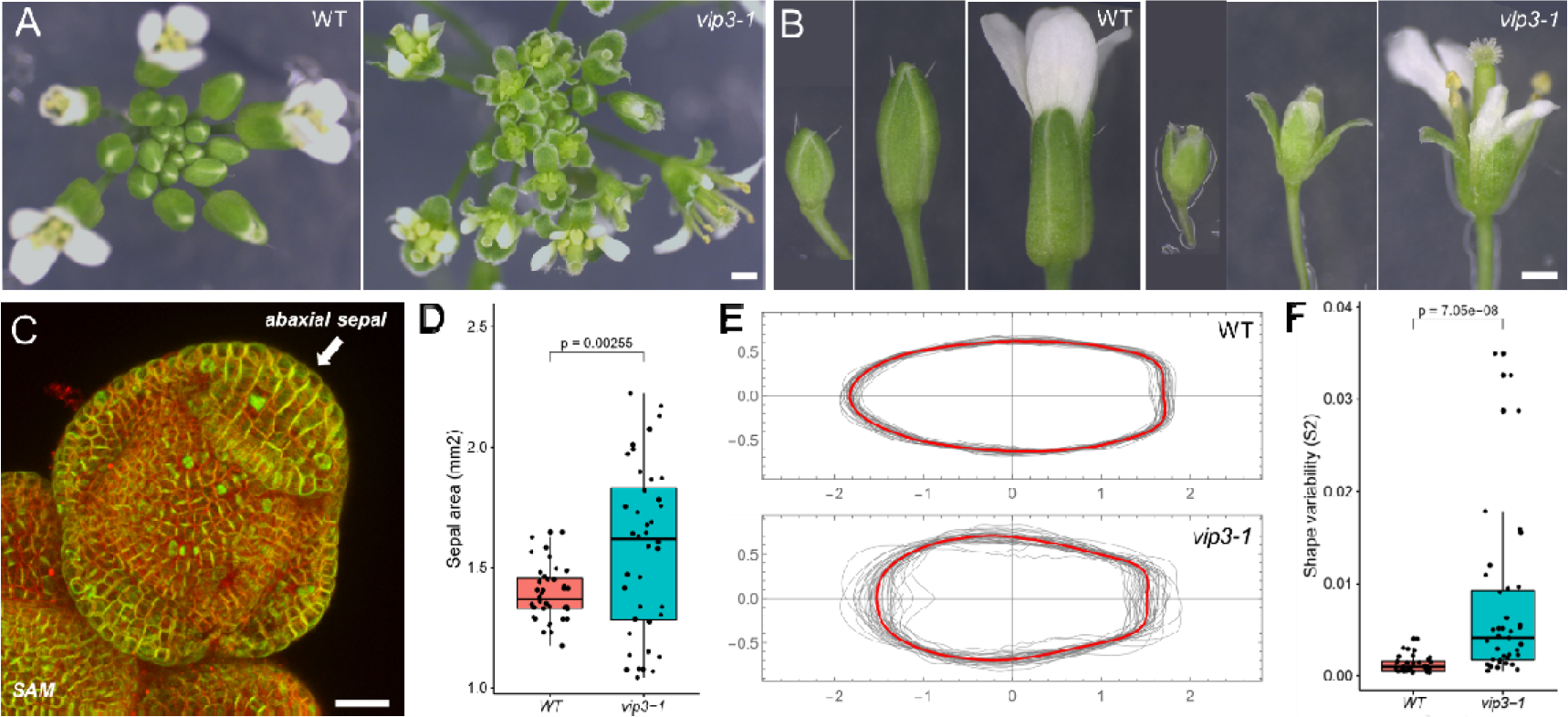
*vip3-1* sepals are more variable in shape and size (A) Top-view of WT and *vip3-1* inflorescences showing the early opening of floral buds and outward-curing of the sepals in the mutant. Scale bar = 1 mm. (B) Close-up images of WT and *vip3-1* floral buds of three different stages. Scale bar = 1 mm. (C) Confocal image of a young WT floral bud to indicate the position of the abaxial sepal (*i.e.* the sepal farther away from the SAM). Abaxial sepals are used throughout the present study. Scale bar = 20 µm. (D) Size analysis: quantification of WT and *vip3-1* abaxial sepal area (mm^2^). *vip3-1* abaxial sepals are bigger and more variable in size. (E) Shape analysis: plots showing the contours of WT and *vip3-1* abaxial sepals, after normalization to the area. The red outline is the average shape. *vip3-1* sepals are more variable in shape compared to the WT. (F) Quantification of Sepal shape variability *S_2_* (squared deviation of sepal outlines) confirming the higher variability in shape in *vip3-1*.

Regarding sepal size, WT flowers produced abaxial sepals of remarkably comparable area (mean ± SD 1.39 ± 0.11 mm^2^; Figure 2D, n_WT_= 40 sepals). In contrast, the *vip3-1* mutant produced abaxial sepals of slightly bigger but highly variable area (mean ± SD 1.57 ± 0.35 mm^2^; Figure 2D, n*_vip3-1_*= 40 sepals). We found the coefficient of variation for *vip3-1* sepal area to be 3 times larger than that of the WT (CoV_WT_ = 7.8%, CoV*_vip3-1_* = 22.2%).

To analyze sepal shape variability in *vip3*, we next focused on the geometry of the sepal contour. To account for size variability in the mutant, the contours of all sepals were extracted and normalized by their area to make the analysis independent of organ size, and then plotted (Figure 2E). Quantification of variability in sepal contour indicated that the median shape variability *S_2_* for *vip3-1* abaxial sepals was about 4 times larger than that of WT (Figure 2F; median ± SE 0.98 ± 0.14 10^-3^ for WT *vs.* 4.1 ± 1.37 10^-3^ for *vip3-1*, *p* =7.05e^-8^, Kruskal–Wallis test). Altogether, these analyses indicate that *VIP3* loss of function leads to increased variability in sepal shape and size.

### Growth in the *vip3* abaxial sepal is locally more heterogeneous

The analyses above showed that in *vip3-1* local variability of gene expression correlates with increased shape variability in abaxial sepals. In the simplest scenario, variability in gene expression would increase cell growth variability, possibly leading to final organ shape variability. To test that scenario, we next quantify cell growth in developing abaxial sepals.

Mature abaxial sepals in *vip3-1* are more variable in shape and size, yet abaxial primordia in both backgrounds initiate with comparable size variability (Figure S2A-B). This further confirms that the variable abaxial sepal size and shape in *vip3-1* relate to sepal growth, and not to defects in floral patterning or early sepal initiation (at least when selecting *vip3-1* flowers with 4 sepals).

To check how growth contributes to variable sepals in the *vip3-1* mutant, we tracked cell growth of WT and *vip3-1* abaxial sepals over a period of 7 days. Abaxial sepals expressing the membrane marker *pUBQ10:Lti6b-TdTomato* were imaged every 24 hours using confocal microscopy. At the first time point, they were around stage 4 according to (Smyth et al., 1990). Confocal stacks were then denoised using the AniFilters tool described in (Long et al., 2020) and imported to MorphoGraphX (de Reuille et al., 2015) for cell segmentation and growth analyses. Representative growth map for WT and *vip3-1* sepals over the period is shown in Figure 3A-B. Here we focus on growth heterogeneity between neighboring cells. We will describe the larger growth kinetics and pattern in more details in the next part.

**Figure 3.**
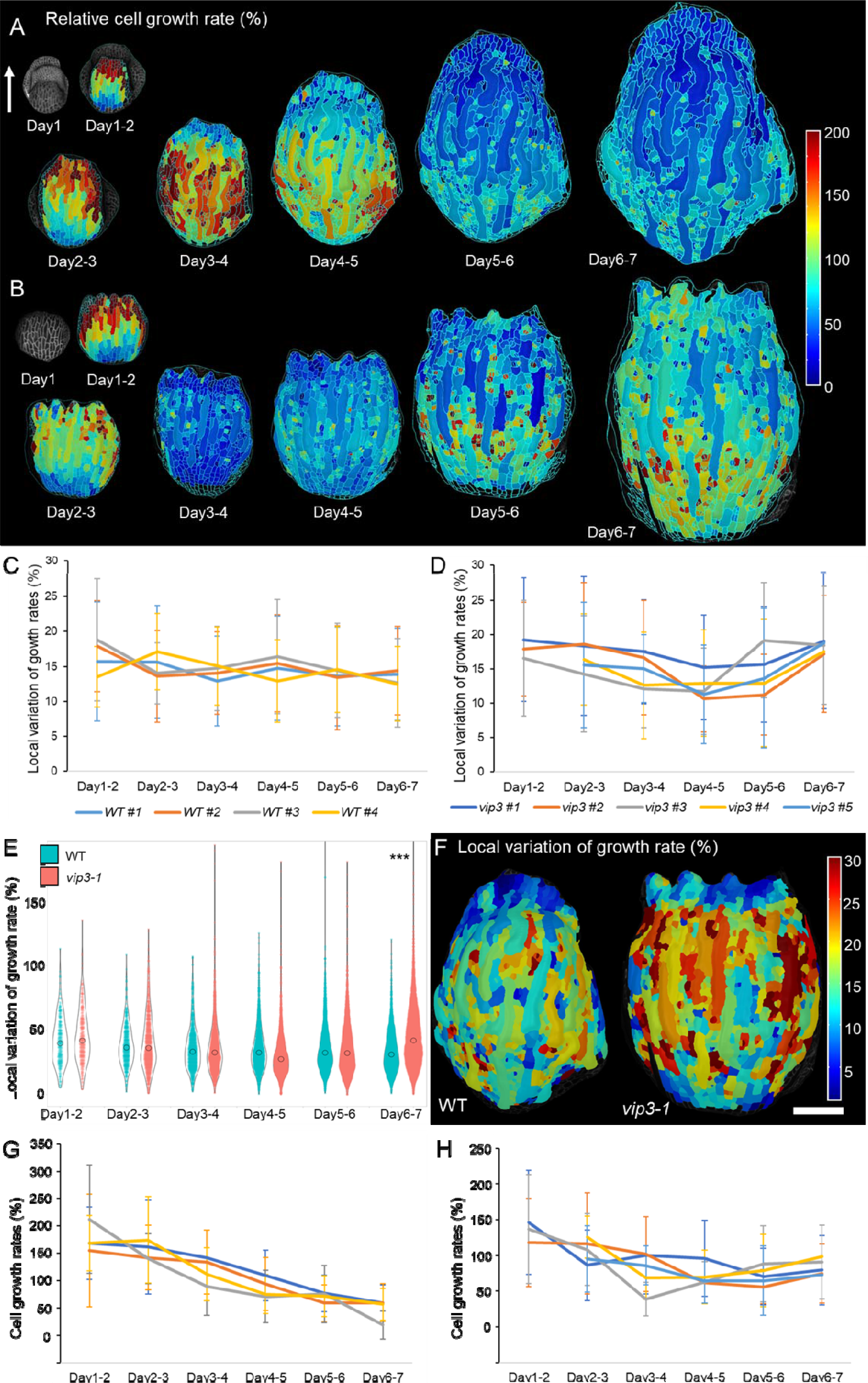
Growth analysis of *vip3-1* sepals over seven days reveals increased growth heterogeneity and a failure to arrest growth at later stage. (A-B) Heat map of cell growth rate (in area) over 24-h intervals of WT and *vip3-1* sepals, projected on the second time point (*e.g.* growth rate of a sepal from Day1 to Day2 is projected on the Day2 sepal). The arrow indicates the main axis of the sepal, pointing to the tip. The same scale and magnification are used. Scale bar = 100 µm. (C-D) Local variation of growth rates (%) of individual WT and *vip3-1* samples. Note that WT samples show fluctuations around a stable level, but *vip3-1* do not. (E) Summary of local variation of growth rates from individual samples in (C-D). Kolmogorov-Smirnov test confirms that CoV of growth rates in WT and *vip3-1* are from two different distributions; *p_WT vs vip3-1(day6-7)_ = 2.2e^-16^*. (F) Representative heat map showing local variation of growth rates in WT and *vip3-1* sepals at Day6-7 where the variation differs the most between two genotypes. Scale bar = 100 µm. (G-H) Cell growth rates over 24-h intervals of individual WT and *vip3-1* sepals. While WT samples exhibit a declining growth rate over time, *vip3-1* sepals maintain and even increase their growth at late stages. For averaged trends, see Figure S3.

Local growth heterogeneity here is defined as coefficient of variation (CoV) in growth rates of cells within a group; the group is formed by a cell and all other neighbors contacting it (Uyttewaal et al., 2012). From the sepals in the time-series experiment described above, we used MorphoGraphX to find all the groups and calculate their local growth heterogeneity, expressed as CoV (%) of local growth rates. Data for individual WT and *vip3-1* samples are shown in Figure 3C and 3D respectively, while Figure 3E shows the grand average of Figure 3C and 3D.

We found that during the period, local growth heterogeneity in WT sepals was surprisingly stable: the average CoV of local growth rates for each sample fluctuate around 14.7% during the 7-day period (Figure 3C). Individual samples showed slight fluctuation in local growth rates between days. This is remarkable, knowing that average cell growth rate in the WT steadily reduces from ∼176% to ∼49% within the same period (Figure 3A and 3G).

In *vip3-1* sepals, we observed more fluctuations in local growth heterogeneity between time points (Figure 3D). More specifically, it went down from ∼17.9% to ∼12.2% in the first four days, then quickly increased to ∼18.2% in the last three days (Figure 3E). Such increased variability was also observed when considering the growth history of individual samples: kinetics of local growth heterogeneity in the *vip3-1* sepals were more erratic than those in the WT (Figure 3C and 3D).

We then mapped local growth heterogeneity onto the sepal meshes, with a focus on the last time point where the mutant shows significantly higher growth heterogeneity (CoV = 13.4% for WT vs. 18.2% for *vip3-1*). Figure 3F illustrates how local growth heterogeneity is more intense and widespread in *vip3-1*, when compared to the WT.

The increase in local growth heterogeneity also happened in another developmental context. We performed the same analyses on SAMs growing over a 24h period and found that cells in *vip3-1* SAMs also displayed higher local variation of growth rates (Figure S3A-B). Regarding stomatal guard cells, we found that the difference in size between two sister guard cells is significantly higher in the mutant when compared to WT counterparts (median difference = 7.3% for WT, = 12.5% for *vip3-1*, *p =* 3.2*e^-6^* Kruskal–Wallis test, n_WT_ = 171, n*_vip3-1_* = 169 pairs) (Figure S3C). Although this is a snapshot of mature stomata instead of a growth kinetics, it suggests a that growth difference between two sister guard cells in the mutant is higher compared to those in the WT. To conclude, these analyses reveal an increased heterogeneity in growth rates between adjacent cells in *vip3-1*.

### Impaired boundaries between growth domains in *vip3*

So far, we have shown that increased local variability in gene expression correlates with increased local growth heterogeneity in *vip3-1*. Yet, this does not explain how this could translate into variable organ shape. In particular, many compensatory mechanisms have been previously shown to regulate organ development (Horiguchi & Tsukaya, 2011). To reveal the nexus between local growth heterogeneity and final organ shape variability, we next analyzed global growth patterns in WT and *vip3-1* sepals.

First, we quantified average growth rates of WT and *vip3-1* sepals. Growth kinetics of WT sepals were relatively similar to one another (Figure 3G). In the first four days (Day1-4) the overall growth rate gradually decreased (from 176% in Day1-2 to 119% in Day3-4, *i.e.* 32% reduction), and it decreased faster afterwards. Overall, *vip3-1* sepals also displayed a growth trend in most of the experiment period (Figure 3H). Compared to the WT, growth rates of *vip3-1* sepals decreased more quickly in the first four days (from 134 % in Day1-2 to 79% in Day3-4, 41% reduction). More surprisingly, we observed that sepal growth rates then stopped decreasing and eventually increased between Day4 and Day7 in the mutant. The contrasting behaviors in growth between WT and *vip3-1* sepals are illustrated in Figure S3D which displays the average sepal growth rates of the two genotypes. Note that whereas all analyzed WT sepals displayed rather consistent growth kinetics, growth kinetics in *vip3-1* sepals were also more variable (Figure 3G and 3H).

Next, we analyzed the differences in regional growth patterns between the two genotypes. Figure 3A shows a representative heat map of the growth pattern in WT sepals. WT sepals exhibited the typical growth dynamics as already described in (Hervieux et al., 2016). In the first three days, sepals displayed a sharp growth gradient from the tip to the base, with the tip growing much faster than the rest of the sepal. From Day3 onward, growth rate at the tip plummeted and the maximum of growth rate shifted to cells in the middle and later on at the base of sepals.

v*ip3-1* sepals displayed a similar growth pattern in the first days compared to WT ones, with the tip growing faster than the rest of the sepals (Figure 3B). However, the difference between the slow-growing area and fast-growing area was less prominent in the first five days (Day1-5) when compared to the WT. Interestingly, in the last two days, cells at the tips of *vip3-1* sepals reactivated their growth, while WT tips grew very slowly.

Thus, not only does *vip3-1* exhibit higher local growth variability between adjacent cells, it also exhibits a noisier growth pattern at global scale. More interestingly, we also found that *vip3-1* is unable to generate a sharp growth arrest front at the sepal tip, as observed in the WT. The sharp difference between slow-growing tip and fast-growing middle area in the WT sepal (Day3-5 in Figure 3A) has been proposed to act as a mechanical signal for growth arrest in the sepal. In short, such a mechanical conflict prescribes a transverse tensile stress pattern, to which cells resist by reinforcing their cell walls, ultimately leading to growth arrest at the tip (Hervieux et al., 2016) (see also Figure 4D). The increased local growth variability in *vip3-1* may prevent the build-up of such a regional conflict. This scenario is also consistent with the presence of a wider tip in *vip3-1* sepals, mimicking mutants with reduced response to mechanical stress at the sepal tip (Hervieux et al., 2016).

**Figure 4.**
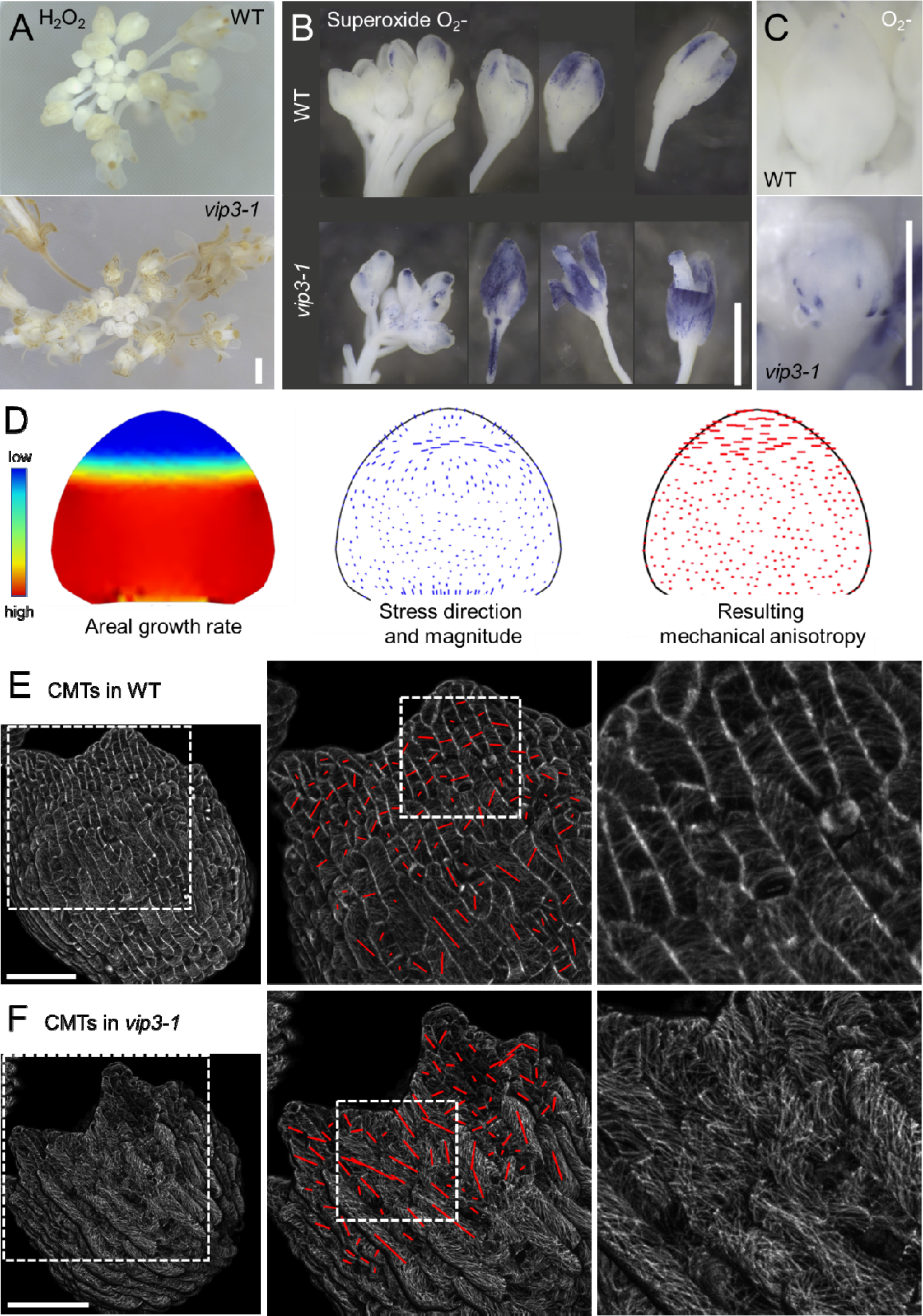
Patchy ROS production and noisy MT alignment in *vip3-1* sepals. (A) DAB staining for H_2_O_2_ in WT and *vip3-1* inflorescences. Note the increased staining in *vip3-1*. Scale bar: 1mm. (B) NBT staining for superoxide in WT and *vip3-1* inflorescences. Note the increased staining in *vip3-1*, as well as the patchy pattern in young *vip3-1* sepals. Scale bar: 1mm. (C) NBT staining for superoxide in young flower buds (close-ups). Note the patchy staining in *vip3-1*, but not in WT. Scale bar: 1mm. (D) Differential growth between the center domain and the tip domain of the sepal prescribes a maximal transverse tensile stress direction, with which cortical microtubules align. The resulting cellulose- dependent mechanical anisotropy in the wall resist stress, and contributes to growth arrest (reproduced from (Hervieux et al., 2016)). (E) Organization of cortical microtubules (CMTs) in WT sepals. The transverse CMT pattern at the tip matches the predicted supracellular maximal tensile stress direction shown in panel D. Scale bar: 50 µm. (F) Organization of cortical microtubules (CMT) in *vip3-1* sepals. In contrast to the WT, *vip3-1* sepals do not display a consistent transverse CMT band at the tip. Scale bar: 50 µm.

### ROS production and MT alignment reflects diffuse mechanical conflicts in *vip3*

If mechanical conflicts are more distributed in the *vip3*-1 mutant, this should be reflected in the cell response to stress in the *vip3-1* sepals. Two classic markers of such mechanical stress response are reactive oxygen species (ROS) production and cortical microtubule (CMT) alignment.

ROS accumulates in sepals from tip to base coincident with their growth gradient, with a maximum at the tip where the mechanical conflict occurs (Hong et al., 2016). We examined ROS levels in WT and *vip3-1* sepals using chemical stains specific for hydrogen peroxide (H_2_O_2_) and the superoxide radical (O_2_^-^), two major ROS molecules. We found that the levels of both species were much higher in *vip3-1* sepals than in WT ones (Figure 4A for H_2_O_2_ staining, and Figure 4B for O_2_- staining, see also Figure S4A). Interestingly, *vip3-1* sepals exhibited a patchy ROS pattern (O_2_^-^) in very young buds, in contrast to WT floral buds which showed no staining (Figure 4B-C, Figure S4A). We also observed patchy ROS staining in the mutant carpels but not in the WT ones (Figure S4A). These data area consistent with mechanical conflicts distributed more locally in the mutant.

ROS signatures, however, can also reflect different types of stress*, i.e.* beyond mechanical conflicts. To challenge our finding, we thus turned to the organization of cortical microtubules (CMTs), the directional behavior of which can be related to maximal tensile stress directions (Hamant et al., 2008). In the WT, at the stage where differential growth between the tip and the middle of the sepal generates a mechanical conflict (Figure 4D), we could observe a band of transverse CMTs at the boundary between the two domains, as previously shown (Figure 4E, (Hervieux et al., 2016)). In the *vip3-1* mutant, CMTs appeared as thicker bundles, mimicking cells experiencing higher level of stress (see *e.g.*(Heisler et al., 2010)). More interestingly, the consistent multicellular transverse band of CMT arrays was largely disrupted: most cells exhibited disorganized CMT bundles instead (Figure 4F, Figure S4B).

Altogether, this provides a scenario in which local variability in gene expression in *vip3-1* leads to increased local heterogeneity in cell growth. This prevents the formation of distinct regional growth domains. The resulting mechanical conflicts between domains are thus hindered, and cells keep growing without a constraining factor. In the end, organ shape becomes more variable.

However, while differences in local growth heterogeneity are significant between WT and *vip3*, they might not be sufficient to explain the variability in final organ shape on their own. Other parameters (*e.g.* disrupted hormone patterns) may contribute to organ shape variability in *vip3*. To assess these other contributions in a more integrative way, we next measured the stiffness of *vip3-1* cell walls, reasoning that all growth factors would affect this parameter in the end.

### Reduced cell wall stiffness in *vip3-1* sepals

First, we measured cell wall stiffness using AFM, *i.e.* by indenting the cell wall with a fine tip, and recording the deformation of the cantilever. Through the indentation, cell wall stiffness can be calculated and usually reported as apparent elastic modulus in mega Pascal (MPa). As often for this type of measurements, elastic moduli can be quite variable. First, we worked with sepals of ∼ stage 6 when the bud is fully closed (Smyth et al., 1990) and cuticular ridges are not formed yet (Hong et al., 2017) because the ridges can interfere with AFM indentation. AFM scanning revealed that cells in WT sepals had stiffer cell walls than those in *vip3-1* (Figure 5A). Average apparent elastic modulus of WT cell walls was around 32 MPa, while that in the mutant was about 24% lower at around 24 MPa (*p=*0.00451, *n*_WT_ =6 sepals, *n_vip3-1_* =8 sepals) (Figure 5B).

**Figure 5.**
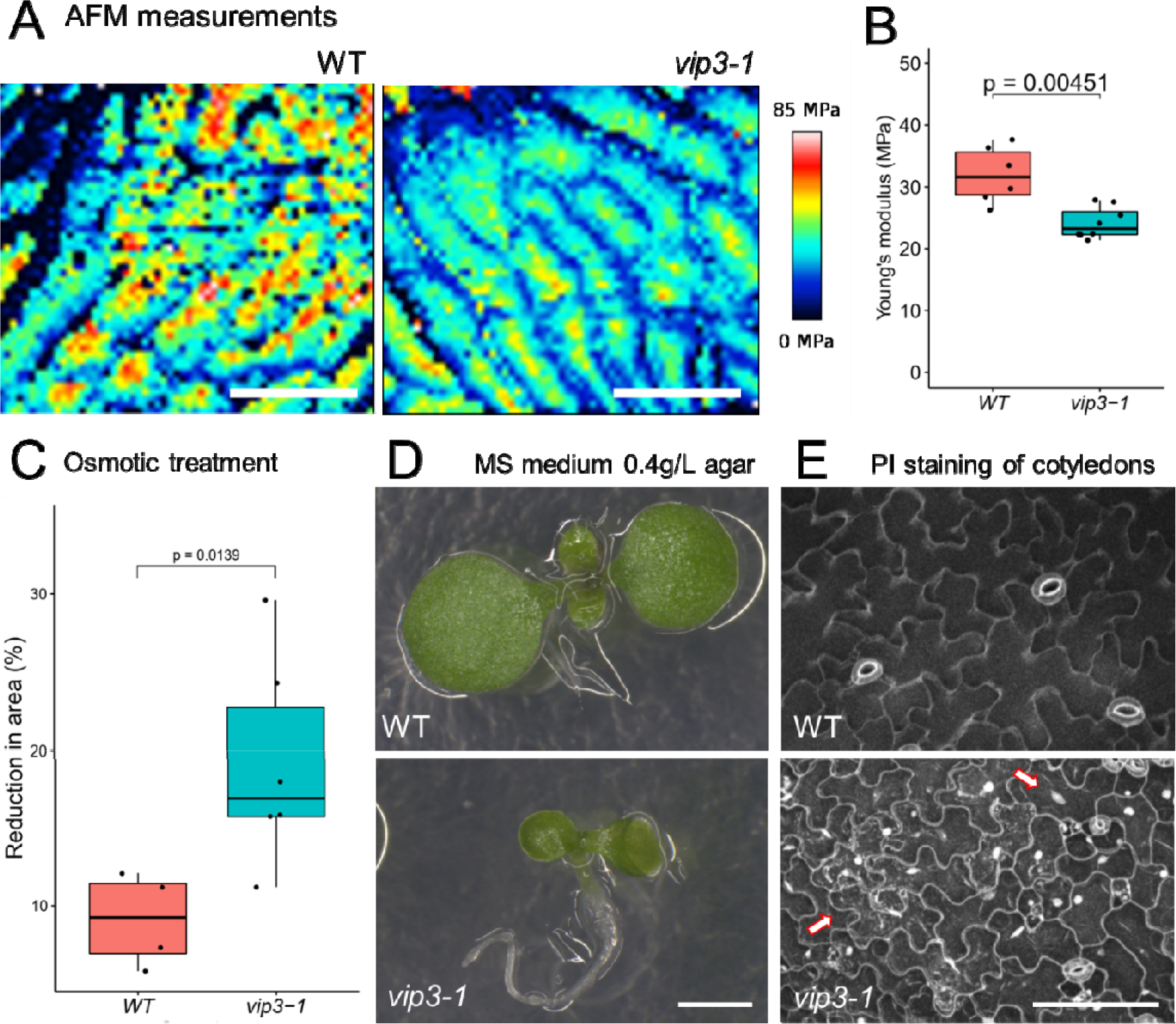
Weaker cell walls in *vip3-1* (A) AFM map revealing cell wall stiffness in WT and *vip3-1* stage 6 sepals. (B) Quantification of apparent elastic modulus (MPa) with AFM. (C) Reduction of sepal area after 30 min of hyperosmotic treatment (NaCl solution). (D) Phenotype of WT and *vip3-1* seedlings growing on MS medium supplemented with 0.4g/L agar. *n*_WT_ =10, *n_vip3-1_* = 10 seedlings. Scale bar = 1mm. (E) Confocal images of WT and *vip3-1* cotyledons stained with propidium iodide (PI) to check for cell viability. Examples of dead cells in *vip3-1* are indicated by the arrows. Scale bar = 100 µm.

To further confirm these measurements, we analyzed the mechanics of sepals using a different approach. The osmotic treatment allows one to assess cell wall stiffness over the whole sepals, while also measuring wall properties in all direction (and not only along the vertical axis of the AFM cantilever (Hong et al., 2016; Kierzkowski et al., 2012). We used NaCl 0.4M solution to induce a hyper-osmotic stress and recorded the deformation of the sepals using MorphoGraphX (see Material and methods). After 30 min treatment, cell area in WT sepals was reduced by 9.1 ± 1.5% (mean ± SE), while in the *vip3-1* sepals the figure was 19.1 ± 2.7% (mean ± SE) (Figure 5C). Altogether, these experiments demonstrate that *vip3* sepals are softer than WT ones.

To further confirm this conclusion, and extend it to other *vip3-1* tissues, we tested the ability of cells and cell walls to resist hypo-osmotic stress (Malivert et al., 2021; Verger et al., 2018). To test this hypothesis, we subjected the mutant to such conditions, using MS medium containing low concentration of plant agar (0.4% agar instead of 0.8% in normal MS). Medium with low agar concentration has higher water potential, and this high water potential increases tensile stress in cell walls (Verger et al., 2018). In normal medium (0.8% agar), *vip3-1* seedlings could grow relatively well, although they were smaller than WT (Figure S5A-B). However, in 0.4% agar medium, *vip3-1* seedlings were much smaller than WT ones, and displayed signs of necrosis at 5 days after sowing (Figure 5D). To check for the presence of dead or dying cells, *vip3-1* seedlings were stained with propidium iodide (PI) which cannot penetrate plasma membrane of healthy cells and stains the nuclei of dead cells. Many dead cells or clusters of dead cells in the *vip3-1* cotyledons were observed (Figure 5E, arrows point at stained nuclei and other intracellular staining). No dead cells were observed in WT cotyledons in the same conditions (Figure 5E). We conclude that weaker cell walls in *vip3-1* could not bear the high osmotic pressure in the cells when the mutant grew in high water potential medium, leading to cell bursting.

Altogether, AFM and osmotic treatments point to softer cell walls in *vip3-1*, in sepals as well as in the whole plant. This is also consistent with the observed increase in overall ROS accumulation and CMT bundling.

### High organ variability is not due to weaker cell walls

So far, we have shown that *vip3-1* mutant display much higher abaxial shape variability (Figure 2E-F), and that *vip3-1* has weaker cell walls in general. The question now is, could weaker cell walls be sufficient for sepal shape variability? In other words, do plants having weaker cell walls display high sepal variability like the *vip3-1* mutant? Plant cell wall is made of load-bearing cellulose microfibrils that are tethered by a matrix of hemicelluloses (xyloglucans and xylans) and pectins (Lampugnani et al., 2018). For our purpose, we selected mutants of *KATANIN, XXT1 XXT2 (XYLOSYLTRANSFERASEAND1* and *2*), and *FERONIA*. KATANIN is involved in cortical microtubule organization and consequently cellulose biosynthesis (Burk et al., 2001), while XXTs are involved in xyloglucan biosynthesis (Cavalier et al., 2008). FERONIA is a receptor kinase involved in maintaining cell wall integrity (Feng et al., 2018; Malivert et al., 2021). There is ample evidence demonstrating that both *katanin (bot1-7)* and *xxt1 xxt2* mutants have significantly weaker cell walls than the WT in differentiating tissues (Bichet et al., 2001; Burk et al., 2001; Burk & Ye, 2002; Cavalier et al., 2008; Park & Cosgrove, 2012; Ryden et al., 2003). *feronia* mutant (*fer-4*) displays cell death when growing in MS medium containing low concentration of plant agar (high water potential), comparable to the response of *vip3-1* (Malivert et al., 2021). We quantified sepal shape variability (*S_2_*) in mature sepals using the SepalContour tool. We found that sepal shape variability in *bot1-7*, *xxt1xxt2* and *fer-4* is not significantly higher when compared to the WT, and significantly lower when compared to *vip3-1* (Figure S5C-D) (median ± SE 1.15 ± 0.32 10^-3^ for WT, 1.25 ± 0.39 10^-3^ for *bot1-7,* 1.18 ± 0.31 10^-3^ for *xxt1 xxt2*, 1.50 ± 0.17, 2.91 ± 2.90 10^-3^ for *vip3-1*, Kruskal–Wallis test). This demonstrates that high sepal shape variability in *vip3-1* is not simply due to weaker cell walls.

### Softer cell wall enhances the expression of local mechanical conflicts in *vip3*

We now have a refined model in which local variability in gene expression together with the presence of softer walls leads to strong mechanical conflicts between adjacent cells growing at different rates. This hinders the formation of sharp regional boundaries and reproducible organ shapes. If this scenario were true, we should be able to detect evidence of even stronger local conflicts between adjacent cells when cell ability to respond to mechanical stress is reduced.

To test this hypothesis, we decreased even further the ability of cells to reinforce their cell walls by applying oryzalin, a microtubule depolymerizing drug. WT and *vip3-1* sepals were treated with 20 µg/ml oryzalin for 3hr every 24 hours, and samples were imaged right before each oryzalin treatment, at t=24h and t=48h (see Figure S6A for the experiment plan). We choose to analyze the impact of depolymerizing microtubules on growth between adjacent cells at Day2-Day3 where the impact of oryzalin is clear (Figure S6B). Oryzalin-treated sepals in both genotypes kept growing during the 3-day period, even though cell division is prohibited. In this condition, *vip3-1* sepals also display a higher local variation of growth rates (CoV_WT_ = 16.8% vs. CoV*_vip3-1_* = 20.8%, Figure 6A-B and Figure S6C). More interestingly, we observed that while some small cells (as view from the top, surface area < 30 µm^2^) in the WT did not grow much (Figure 6C, arrow), some small cells in *vip3-1* exhibited negative growth, *i.e.* were contracted by their neighbours (Figure 6D, arrows). Changes in cell surface area over the two days of small cells of similar size in WT and *vip3-1* sepals are illustrated in Figure 6E. Orthogonal views of the squeezed cells in *vip3-1* (along the red lines in Figure 6D) showed that the spaces between anticlinal walls of these cells did become smaller, even disappeared (Figure 6F and Figure S6D-E). 3D reconstruction and growth measurement of these cells confirmed their volume contraction and the disappearance of a cell (arrows, Figure 6G-I). We could not find such strong growth conflict in the WT.

**Figure 6.**
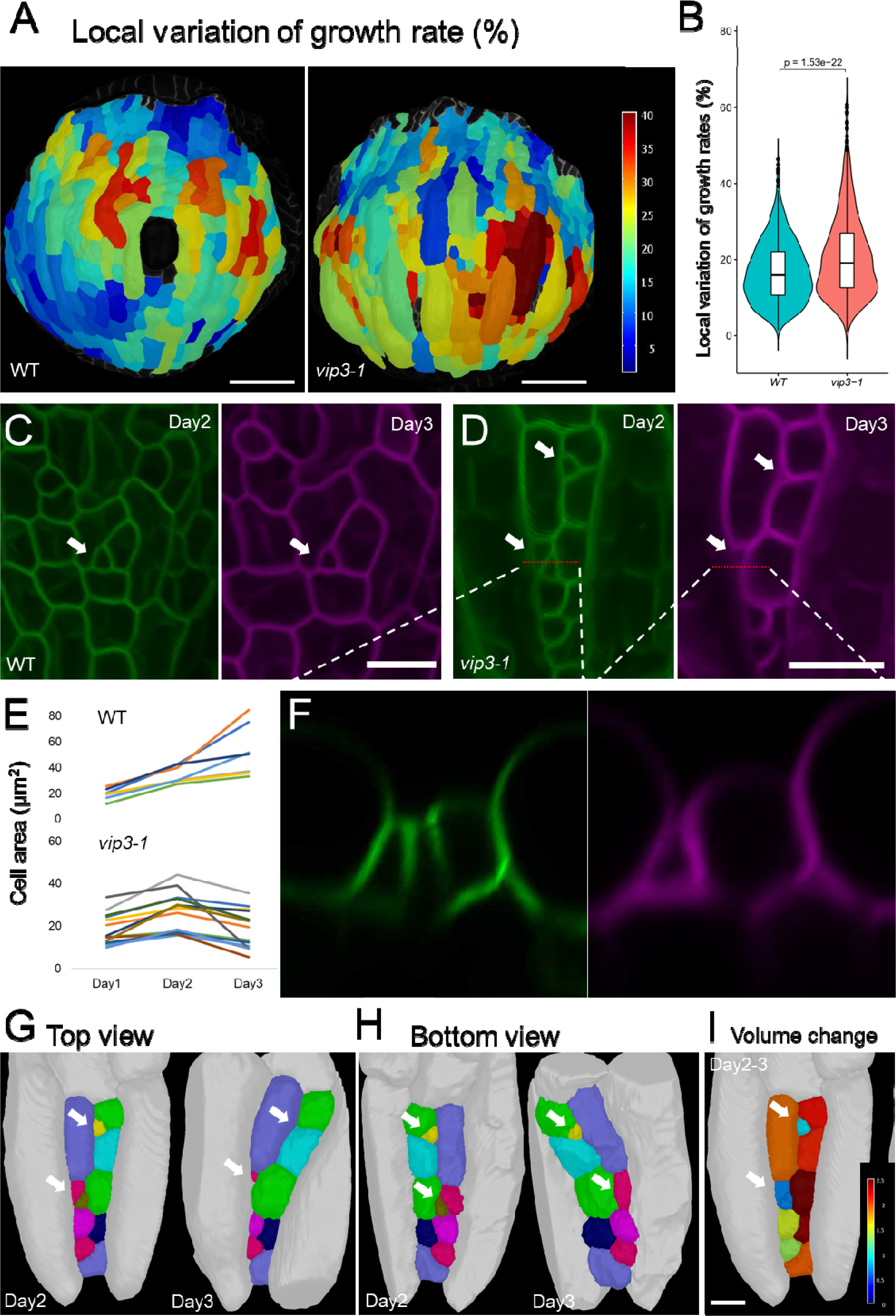
Oryzalin-induced microtubule depolymerization amplifies local mechanical conflicts in *vip3-1* (A-B) Local variation of growth rates (%) in WT and *vip3-1* sepals treated with oryzalin for 2 days. (C-D) Changes in cell surface area of small cells in WT and *vip3-1* over time (arrows). (E) Growth kinetics (in cell surface area) of small cells in WT and *vip3-1*. Note the reduction in surface area in *vip3-1* cells, which is not seen in the WT. Each line represents a cell. *n*_WT_ = 9 cells, *n_vip3-1_* = 13 cells. (F) Orthogonal views of the squeezed cells in *vip3-1* along the red lines showed in Figure D. Note the reduction or disappearance of the space between anticlinal walls. See also Figure S6D-E. (G) 3D reconstruction of some selected cells in (D) at Day2 and Day3, top view. The same cell at Day2 and Day3 has the same color. Note the reduction in surface as viewed from the top of the cells indicated by the arrows (the same cells in (D)); the brown cell at Day2 cannot be detected anymore at Day3. (H) The same cells in (G), bottom view. Note that the brown cell is disappeared in this view as well. (I) Heatmap showing changes in cell volume from Day2 to Day3, projected to Day2. The scale is from 0-2.5 (times). Volume change < 1 means a volume contraction, which happens to the cells indicated by the arrows. The two cells indicated by the lower arrow are counted as one for this measurement.

Altogether, this is consistent with a scenario in which transcriptional noise scales mechanical conflicts down to adjacent cells (even amenable to generate a novel form of cell competition), thereby hindering the formation of sharp boundaries between domains, and ultimately impairing growth arrest mechanisms and leading to variable organ shapes.

## DISCUSSIONS

In this study, we find that Paf1C reduces transcriptional noise in a multicellular context, and we provide evidence for the requirement of noise management to allow mechanical conflicts to develop between regions, instead of between adjacent cells. We propose that channeling growth heterogeneity to a regional scale fuels organ shape reproducibility (Figure 7).

**Figure 7.**
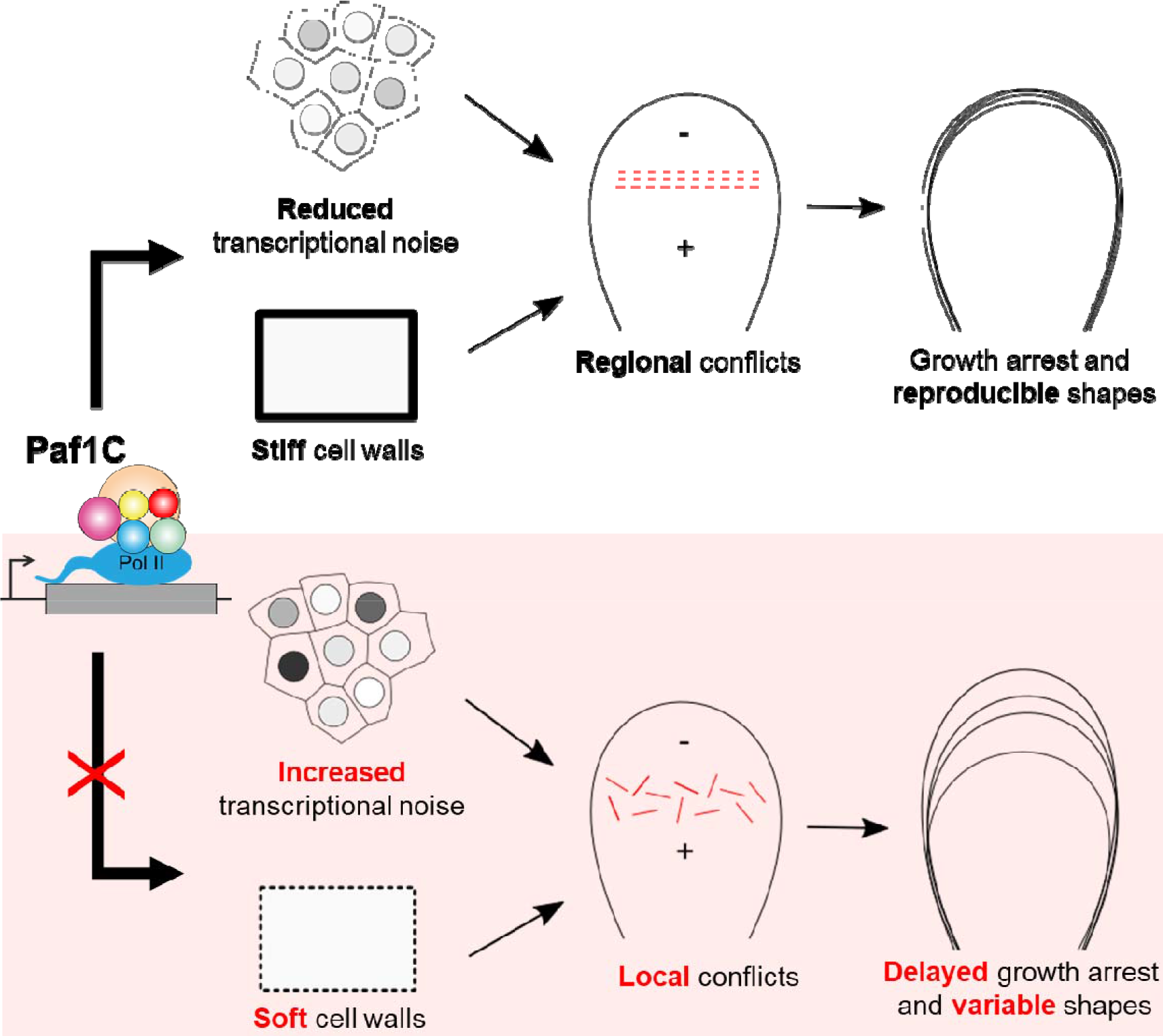
A multiscale model relating Paf1C-dependent transcriptional control to organ shape robustness. Paf1C-dependent transcription contributes to cell wall synthesis and stiffening, while also channeling gene expression variability. Reduced local gene expression variability allows the formation of regions with consistent growth rates, and thus sharp boundaries between them. The associated supracellular mechanical conflict acts as a signal for growth arrest, hence contributing to reproducible shapes. In the *vip3-1* mutant where Paf1C function is impaired, transcriptional defects leads to softer cell walls and increased transcriptional noise. Increased local gene expression variability fuels local growth conflicts between adjacent cells, while weaker cell walls enhance average stress levels. Consequently, growth patterns become noisier and growth arrest is delayed, contributing to variable shapes.

### Transcriptional noise in plants is under genetic control

There are several techniques to quantify transcriptional noise in single cells, but a fluorescent protein targeted to the nucleus is the most practical approach for multicellular systems like plants at the moment. Note that studies on transcriptional noise commonly employ the dual reporter strategy to separate intrinsic and extrinsic noise (*e.g.* to see if noise of the two reporters co-vary or not) (Araújo et al., 2017; Elowitz et al., 2002; Raser & O’Shea, 2004). Here we used a single reporter approach which informs us about the noisiness of transcription regardless of sources.

One of the difficulties with working on transcriptional noise in a multicellular system is that each cell has a different history, and thus the observed noise can rather reflect that history. We circumvented this problem by using sister guard cells, for which the origin and history is by definition very close. Working with different reporter lines, we confirmed that there is a certain level of noise in gene expression in WT Arabidopsis, and that the noise levels vary depending on the genes (Araújo et al., 2017; Meyer et al., 2017). This was also validated when considering other tissues, and in particular SAMs and floral buds.

More importantly, we demonstrated that mutation in *VIP3*, a component of Paf1 complex indeed leads to increased transcriptional noise in different contexts. This is in agreement with the pioneer work showing increased transcriptional noise due to mutation in Paf1C subunits in yeast (Ansel et al., 2008). Among the gene targets, we were particularly surprised to see strong local variability in the expression of *STM*, a key regulator of meristem maintenance. This observation opens many new questions: could redundant factors (*KNAT* genes) buffer such variability in part by displaying “expression averaging” at regional scale or instead amplify it locally by adding even more stochasticity? Could certain master regulators of development, like *STM*, be over sensitive to transcriptional regulators, and thus be more likely to be noisy?

There is evidence showing that VIP3 can participate in two independent complexes, Paf1C in the nucleus and SKI (Superkiller) in the cytosol (Dorcey et al., 2012). These complexes control mRNA production and turnover, respectively. However, while the only available mutant of the SKI complex *Atski2* does not produce any flower development phenotype (Dorcey et al., 2012), mutants of different Paf1C subunits show similar flower development defects (Zhang et al., 2003), strongly suggesting that flower development defects in *vip3-1* are the result of compromised Paf1C, not SKI, activity. Nevertheless, here we used *vip3-1* as a background in which transcription is noisier than the WT regardless of the precise mechanisms. With the identification of VIP3, and probably other subunits of the Paf1 complex, as regulators of transcriptional noise, the corresponding mutants now offer a model system to study the effects of increased transcriptional noise on many cell and developmental questions.

### Transcriptional noise fuels local mechanical conflicts, instead of regional ones

Here we studied one of these questions: the nexus between transcriptional noise and organ shape reproducibility. To do so, we focused on sepals, an excellent system to study organ developmental robustness in Arabidopsis due to their highly reproducible shape and size (Roeder, 2021). A first striking result is the observation that local variation of growth rates of cells in WT sepals is remarkably similar across time points, slightly fluctuating around 15%. This suggests that in the WT there may exist a limit for local growth heterogeneity because growth of a given cell is constrained by its contiguous neighbours. This could be a major limiting factor for many other developmental features such as size, aspect-ratio or flatness, and this remains to be studied. In the mutant sepals, we observed a significant increase in growth heterogeneity between adjacent cells at the end of the imaging period, *i.e.* where sepals between WT and *vip3-1* become progressively and increasingly different.

High differences in growth rates between neighboring cells can lead to local mechanical conflicts because they are all glued together by cell walls. While it is not possible to visualize mechanical stress, one can have an indication of such stress thanks to other signs. For example, mechanical stress induces ROS production in both plant and animal cells (Aikawa et al., 2001; Benikhlef et al., 2013). Using ROS staining, we found that in young floral buds while there was no sign of ROS accumulation in the WT, ROS staining appears in a patchy pattern in *vip3-1* sepals. Note that the patchy ROS staining was also observed in other developmental context (carpels). This supports a scenario in which some cells in *vip3-1* experience a higher level of mechanical stress related to local growth conflicts.

However, increased ROS production in *vip3-1* may also suggest that the variable shape of *vip3-1* sepals is simply due to an overall increase in stress (whether mechanical or biochemical) in the organ. While we cannot completely rule out this possibility, we note that ROS staining not only increased but was also more patchy. This is different from observations in the *ftsh4* mutants with increased, but widespread, ROS production. In that case, the larger ROS producing regions were proposed to synchronize larger populations of cells, ultimately leading to variable organ shapes (Hong et al., 2016). Here, we propose a different scenario in which local noise instead prevents the formation of such large and consistent regions. Furthermore, the presence of heterogeneous CMT behavior between adjacent cells further confirms that the *vip3-1* phenotype is not only due to an increase in stress levels: if stress was higher, without noise, CMTs would be both bundled and co-aligned. Whereas CMTs are indeed more bundled in *vip3*, they are not forming a consistent and multicellular transverse band, again consistent with increased local noise.

### A new form of cell competition in plants

Because local differences in growth rates correlates with increased transcriptional noise, this leads us to address the question of the management of growth heterogeneity. Transcriptional noise can be viewed as a promoter of growth heterogeneity, in the simplest scenario. This is observed in stomata and sepals in our study. Conversely, the ability of the cell to resist to mechanical stress in such local conflicts could buffer growth heterogeneity in the end. Thus, the observed growth heterogeneity likely does not reflect the full contribution of transcriptional noise to growth heterogeneity. In other words, if the cells were not able to resist such conflicts, growth heterogeneity should be higher.

In animal systems, increasing growth heterogeneity can lead to a phenomenon called ‘cell competition’, in which slow-growing cells are outcompeted and eventually eliminated by fast-growing cells. In that case, the death of slow-growing cells may be caused by both chemical signals and mechanical compression (Brás-Pereira & Moreno, 2018). Plant cells are different compared to animal cells because they have a stiff cell wall which can sustain a substantial mechanical pressure, and their position is fixed relative to other cells. Yet, cell wall is flexible. It can be loosened to accommodate cell expansion, or selectively reinforced to cope with mechanical stress. In the present study, when sepals are treated with oryzalin to remove CMTs, and thus weaken the ability of the cell to reinforce its cell wall, we found that the growth conflict is further amplified, leading to the drastic reduction in volume in some cells in the mutant. This had not been reported before in plants, except for the very peculiar cell contraction step in the zygote, shortly after fertilization (Kimata et al., 2016). Our observation echoes the ‘cell completion’ phenomenon mentioned above for animal systems, and points to mechanical compression as the primary cause. It further suggests that growth conflicts are not just passively managed by the existence of stiff cell walls, but requires active adjustment of cells, most likely with the help of CMTs.

### Transcriptional noise leads to more variable organs

Local growth heterogeneity has been linked to organ robustness. In Arabidopsis, it is found that the shoot apical meristem maintains a certain level of growth heterogeneity between cells, and reduced growth heterogeneity in the *katanin* mutant has been associated with the loss of the typical dome shape and clear organ boundary (Uyttewaal et al., 2012). Sepal cells also displayed significant growth heterogeneity, and reduced growth heterogeneity was found in a mutant with more variable organ shape (Hong et al., 2016). The two counter-intuitive examples suggest that a certain level of local growth heterogeneity is necessary for organ shape robustness.

Nevertheless, sepals in *vip3-1* mutant have higher local growth heterogeneity and much more variable final shapes and sizes compared to the WT. We observed that during their growth, *vip3-1* sepals did not exhibit a sharp difference in growth rates between the tip and the base. This is of particular interest because this sharp growth gradient can generate a transverse mechanical stress at the boundary between two regions which acts as a signal for growth arrest (Hervieux et al., 2016), suggesting that growth arrest in the mutant is not tightly controlled. Remarkably, we found that cells at *vip3-1* sepal tip resumed growth at some points, in contrast with WT cells. Although it is possible that the resumed growth is an unrelated mechanism to compensate for the slower growth rates observed in first days of *vip3-1* sepals, it is still consistent with a loose control of growth arrest in the mutant sepals.

Altogether, this suggests that organ shape reproducibility requires a balance between growth heterogeneity activators and inhibitors. Interestingly, the corresponding factors can act as activators or inhibitors, independent of their molecular identity, but instead depending on scale and context. For instance, Paf1C-dependent transcriptional noise increases local growth heterogeneity, but reduces regional growth heterogeneity. Similarly, CMT-dependent wall reinforcement can reduce or increase growth heterogeneity depending on the tissue context (Uyttewaal et al., 2012). In the end, the introduction of transcriptional noise as an instructive cue is an invitation to revisit many cell and developmental questions with the lens of multiscale dynamics.

## ACKNOWLEDGEMENTS

We thank Richard S. Smith (Department of Computational and Systems Biology, John Innes Centre, Norwich, UK) for help in using MorphoGraphX. We thank Platim (Simone Bovio and Claire Lionnet) for their help in using AFM and confocal microscopy. We are grateful to our collaborators (Arezki Boudaoud, Adrienne Roeder, Richard Smith and Jan Traas) for their comments and help in this project. This work is supported by the European Research Council (ERC, grant agreement No 101019515, “Musix”), by CEFIPRA (grant 6103-1), and by the French National Research Agency through a European ERA-NET Coordinating Action in Plant Sciences (ERA-CAPS) grant (Grant No. ANR-17-CAPS-0002-01).

## STAR METHODS

### Key resources table

| REAGENT or RESOURCE | SOURCE | IDENTIFIER |
| --- | --- | --- |
| Biological samples |  |  |
| Chemicals, peptides, and recombinant proteins |  |  |
| Oryzalin | Chem Service | 19044-88-3 |
| 3,3'-diaminobenzidine (DAB) | Sigma-Aldrich | 91-95-2 |
| Nitroblue tetrazolium (NBT) | Bio-Rad | 293-33-9 |
| Experimental models: Organisms/strains |  |  |
| Arabidopsis Col-0 |  | N/A |
| *pUBQ10::Lti6b-tdTomato* | (Shapiro et al., 2015) | N/A |
| *p35S::GFP-MBD* | (Hamant et al., 2008) | N/A |
| *pH1.1::H1.1-EGFP* | (She et al., 2013) | N/A |
| *pH1.2::H1.2-EGFP* | (Rutowicz et al., 2015) | N/A |
| *pH1.3::H1.3-EGFP* | (Rutowicz et al., 2015) | N/A |
| *pSTM::CFP-N7* | (Landrein et al., 2015) | N/A |
| *pAG::AG-2xVenus* | (Fal et al., 2019) | N/A |
| *pAtML1::H2B-3xGFP::tRBCS* | This study | N/A |
| Software and algorithms |  |  |
| SurfCut (ImageJ) | (Verger et al., 2018) | N/A |
| FibrilTool (ImageJ) | (Boudaoud et al., 2014) | N/A |
| MorphoGraphX | (de Reuille et al., 2015) | N/A |
| SepalContour (Mathematica) | (Hong et al., 2016) | N/A |
| MorphoLibJ (ImageJ) | (Legland et al., 2016) | N/A |
| XPIWIT | (Bartschat et al., 2016) | N/A |

### Plant materials and Growth conditions

Otherwise stated, all experiments were performed on Co-0 ecotype. The *vip3-*1 (Salk_139885) mutant was described in (Fal et al., 2019). The plasma membrane marker line *pUBQ10::Lti6b-tdTomato* was described in (Shapiro et al., 2015), the microtubule reporter line *p35S::GFP-MBD* in (Hamant et al., 2008), *pH1.1::H1.1-EGFP, pH1.2::H1.2-EGFP* in (She et al., 2013), *pH1.3::H1.3-EGFP* in (Rutowicz et al., 2015), *pSTM::CFP-N7* in (Landrein et al., 2015), *pAG:: AG-2xVenus* in (Fal et al., 2019).

For all analyses on sepals, plants were grown on soil at 20°C in short-day conditions (8h light/16h dark) for 3 weeks then transferred to long-day conditions (16h light/8h dark cycle). For work on seedlings, seeds were surface-sterilized using Sodium dichloroisocyanurate (dissolve 88mg in 1.0mL water then add 9mL Ethanol absolute to have the working solution), and placed on squared plates containing Murashige and Skoog (MS) solid medium supplemented with vitamins and plant agar at desired concentrations (8g/L or 4g/L). Plates were kept at 4°C for 2 days and then placed in long-day conditions (16h light/8h dark cycle).

### Construction of AtML1 transcriptional reporter and plant transformation

The transcriptional reporter *pAtML1::H2B-3xGFP::tRBCS* was generated using GreenGate assembly (Lampropoulos et al., 2013). The AtML1 promoter encompasses 3384 bp upstream of transcription start and was previously described in (Schürholz et al., 2018). *Agrobacterium tumefaciens* (Agl-0) based plant transformation was carried out using the floral dip method (Clough & Bent, 1998). Homozygosity was determined by antibiotic resistance (kanamycin) and verification of the fluorescence at the microscope.

### Live imaging of abaxial sepals and shoot apical meristems

Preparation of samples were done as described in (Stanislas et al., 2017), section *Plants grown on soil*. Basically, young main inflorescence stems were cut from the plants expressing *pUBQ10::Lti6b-tdTomato*, then old flower buds were dissected out to reveal young floral buds of desired developmental stages, or to reveal the SAM. Dissected inflorescences were cultured in the apex culture medium descried in (Hamant et al., 2014; Stanislas et al., 2017). The samples were kept in the medium overnight before observation. For sepal growth analyses, abaxial sepals were imaged every 24 hours for 7 days. For big sepals which could not fit within an image acquisition frame, several tiles were stitched together using the Stitching plugin in ImageJ (Preibisch et al., 2009). All confocal images were done using an SP8 laser-scanning confocal microscope (Leica) equipped with a long-distance 25X (NA 0.95) water-dipping objective and a resonant scanner module. For CFP: excitation laser 448 nm, collected range 460-490 nm. For GFP: excitation laser 488 nm, collected range 500-520 nm. For tdTomato protein: excitation laser 552 nm, collected range 570-605 nm.

### Image analysis

For growth analyses, the MorphoGraphX (MGX) 3D image analysis software was used (de Reuille et al., 2015). There are several guides to use the software such as (de Reuille et al., 2015; Stanislas et al., 2017; Strauss et al., 2019). To reveal the general growth patterns of the sepals as depicted in Figure 3, the range of growth rates was set to 1-3 for all time points.

Local variation of growth rate is defined here as the coefficient of variation (CoV = SD/mean*100%) in growth rate within a group of cells consisting of a seed cell in the center and all its contacting neighbors, as described in (Uyttewaal et al., 2012). For a given cell, its neighboring cells be found using the Heat map function in MGX (Heat map type: Walls; Heat map visualization: Geometry; Box Geometry is checked). Local variation of growth rate is calculated for every cell in the sample.

To assess gene expression variability in floral meristems and SAMs, images were taken using confocal microscope (Leica SP8) with z-step of 0.2µm from 2 channel: one for nuclei signal (GFP or Venus), and one for PI staining to detect cell walls. The PI (cell wall) image stack were used for cell segmentation, and each cell is given a label in a cell mesh. For a given cell, its neighboring cells be found using the Heat map function in MGX (Heat map type: Walls; Heat map visualization: Geometry; Box Geometry is checked). A cell and its neighbours form a group which will be used to calculate local variation of signal intensity. The nuclei image stacks were first denoised in MGX and segmented in 3D using XPIWIT (Bartschat et al., 2016). Segmented nuclei were then imported into MGX to create a mesh for each nucleus (cub size (µm): 0.4; smooth passes: 2). The cell mesh (Mesh 1) and nuclei mesh (Mesh 2) were then imported into MGX, and cell labels in Mesh 1 were transferred to nuclei in Mesh 2. By using both the nuclei mesh and original nuclei image, total nuclei signal, nuclei volume and nuclei signal intensity (normalized to volume) were then calculated using Heat map function in MGX (Heat map type: Volume; Heat map visualization: Total signal; Box Geometry and Signal are checked). Because nuclei have the same labels as cells, groups of cells are equal to group of nuclei, coefficient of variation (CoV %) in nuclei signal intensity within a group of nuclei/cells (which signify local variation of signal intensity) could be calculated in the same way as local variation of growth rate.

To assess gene expression variability of reporter lines in a pair of guard cells, confocal images of stomata in seedling cotyledons were acquired in the same way as described above (2 channels, z-step 0.2 µm). ‘SUM slices’ projection was made in Fiji for each stack to have total signals. A ROI is drawn for each nucleus, together with a ROI nearby of identical size to account for background noise. Total signal for each nucleus is measured, minus the background noise. The difference (Δ) in total signal between the two sister guard cells was then calculated, using the nucleus with lower signal as the reference (*i.e.* difference in signal between two sister guard cells divided by lower signal intensity of the two cells and expressed as a percentage).

To visualize and quantify microtubule anisotropy in sepals and hypocotyls, cortical microtubule signal (*35S::GFP-MBD;* 0-5µm from the surface) from confocal images were first extracted using the SurfCut plugin in ImageJ (Verger et al., 2018). Anisotropy of extracted cortical microtubules were then quantified using the FibrilTool plugin in ImageJ (Boudaoud et al., 2014).

### Sepal area measurements

Sepals of stage 14 flowers (Smyth et al., 1990) were dissected and placed as flat as possible on double-sided tape on top of a microscope slide. They were photographed on a black background using a binocular equipped with a camera. A Python program in Linux called SepalContour described in (Hong et al., 2016) were used to extract sepal contours and morphological parameters such as sepal area, length, width, aspect ratio and circularity. Some users may find the installation of the tool difficult; therefore, a complete virtual machine (∼ 9GB in compressed size) with the tool ready to run is available upon request.

### Quantification of sepal shape variability

Sepal contours extracted using the SepalContour tool described above were then used as input for shape analyses. The Contour Analysis program was written in Mathematica and described in (Hong et al., 2016). From the input contours, the program returns a graph of all original contours, a graph of contours normalized by their area and the average contour, and for each sepal contour, an *S_2_* value depicting the squared deviation of that contour from the average contour. These *S_2_* values were used to estimate sepal shape variability within and between genotypes (Hong et al., 2016).

### Measuring stomatal guard cell sizes

Stomatal guard cells taken by confocal microscopy using PI staining were segmented and measured using the MorphoLibJ plugin in ImageJ (Legland et al., 2016).

### AFM measurements

AFM experiments were performed as described and discussed in (Zhao et al., 2019). In brief, we used a silica spherical tip with a nominal radius of 400 nm mounted on a silicon cantilever with a nominal force constant of 42 N/m. The applied force trigger was of 1 µN, corresponding to an indentation of 100–200 nm, used in order to indent the cell wall only (Milani et al., 2011; Tvergaard & Needleman, 2018).

### Osmotic treatment on sepals

Sepals of WT and *vip3-1* mutant were treated with 0.4M NaCl water solution for 30min, and sepal areas before and after treatment were compared using MGX as described in (Hong et al., 2016; Kierzkowski et al., 2012; Sapala & Smith, 2020).

### Detection of Reactive Oxygen Species

*In situ* hydrogen peroxide (H_2_O_2_) and superoxide radical (O_2_^-^) were detected using 3,3’- diaminobenzidine (DAB) and nitroblue tetrazolium (NBT) respectively, as described in (Dutilleul et al., 2003; Hong et al., 2016). After staining, the samples were photographed using a Leica binocular.

### Oryzalin treatment

Oryzalin treatment on sepals was done as described in (Hervieux et al., 2016). Basically, sepals were treated with 20 µg/ml oryzalin for 3hr every 24 hours to deplete microtubules. They were then washed several times with water to remove oryzalin. They were imaged with the SP8 before the treatment. Orthogonal views of sepal cells were made in MorphoGraphX using Clip functions. Measurements of surface area of small cells in oryzalin-treated sepals were done manually using Fiji. Measurements of cell volumes were done in MorphoGraphX using the ITK functions (Stack > ITK > Segmentation > ITK Watershed Auto Seeded).

### Statistical analyses and Data visualization

Statistical analyses were performed in R (R Core Team, 2013). Graphs were created in R using ggpubr (Kassambara, 2020a) and rstatix (Kassambara, 2020b) packages, or the online tool PlotsOfData (Postma & Goedhart, 2019), or in Microsoft Excel.

---

**Figure S1.**
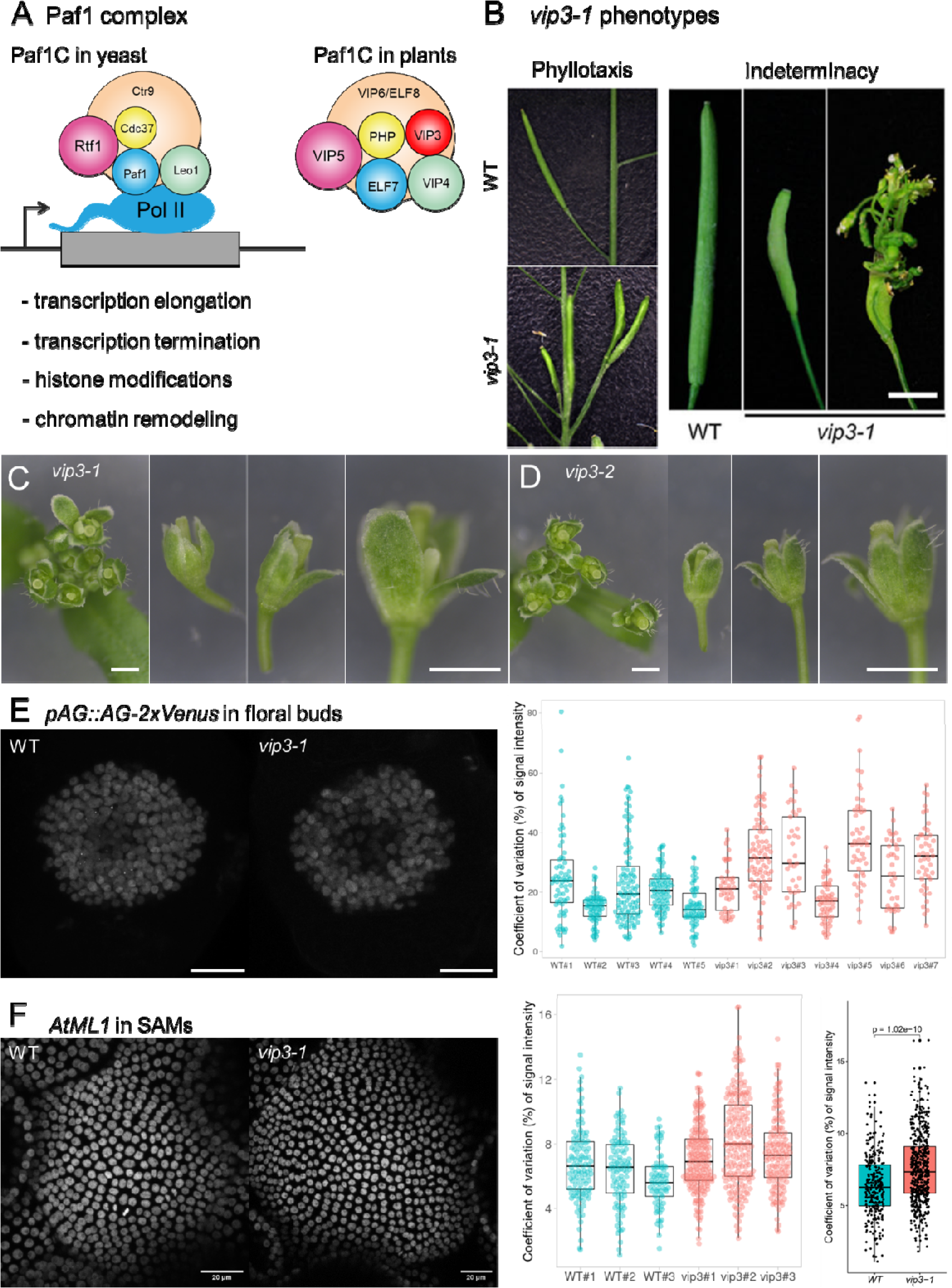
Related to Figure 1. VIP3 as a regulator of transcriptional noise (A) Diagrams showing the components of the Paf1 Complex (Paf1C) in yeast and in plants. The Paf1C binds to the RNA Polymerase II and is involved in controlling transcription elongation, transcription termination as well as histone modifications and chromatin remodelling. Adapted from (Crisucci & Arndt, 2011). (B) Mutation in *VIP3* results in many phenotypes, including defective phyllotaxis and flower indeterminacy in which the flower can produce a whole inflorescence. Adapted from (Fal et al., 2017, 2019). (C,D) Images of *vip3-1* (C) and *vip3-2* (D) inflorescences and young flowers. Scale bars = 1 mm. (E) Representative image of *pAG::AG-2xVenus* expression in stage 3 floral buds (left) and coefficient of variation (%) of signal intensity between neighbouring cells (right) for the WT and *vip3-1* samples. (F) Similar to (E), but for *pAtML1::H2B- 3xGFP::tRBCS* in WT and *vip3-1* SAM samples. All samples are pooled together in the last graph (Kruskal–Wallis test).

**Figure S2.**
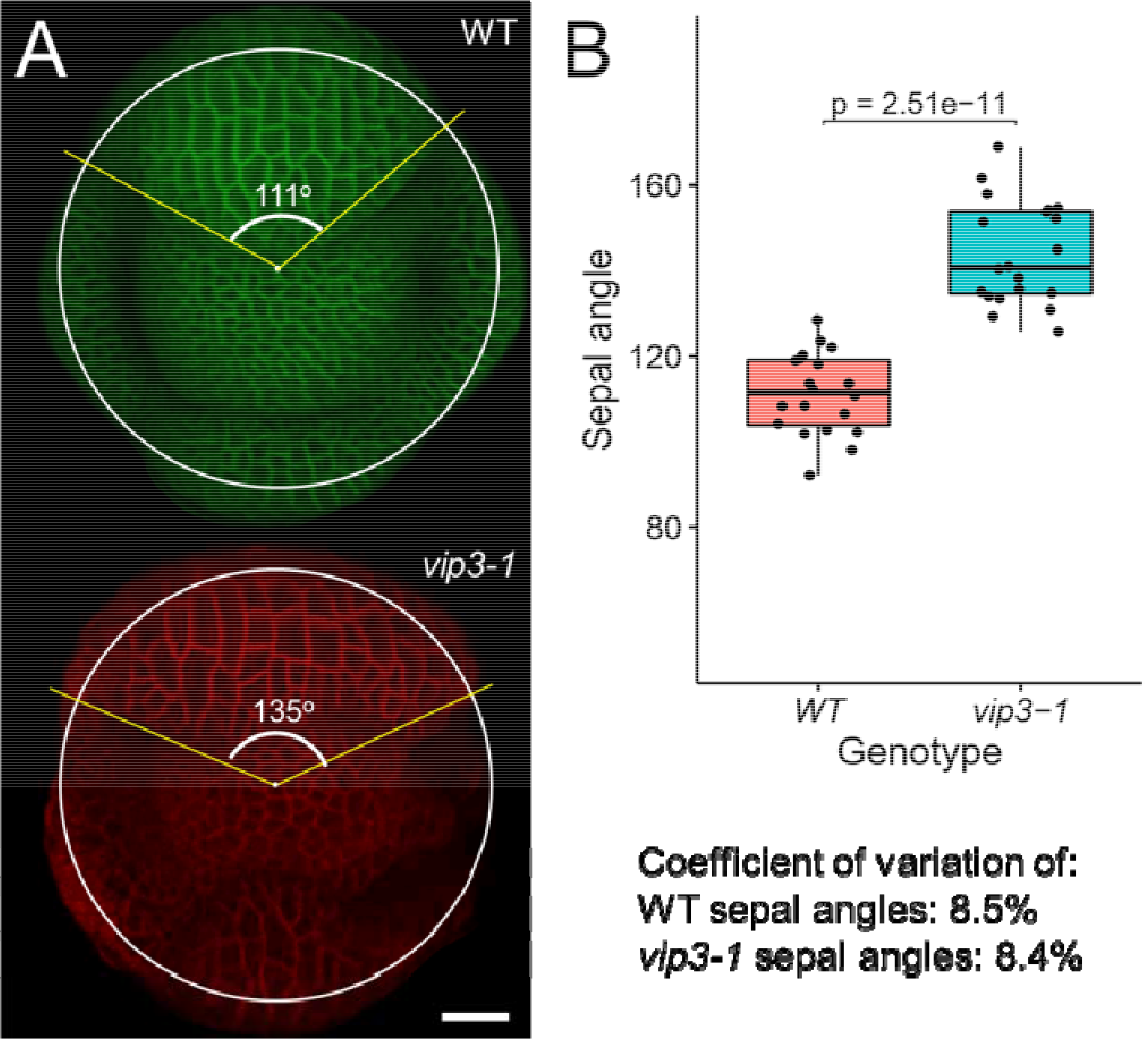
Related to Figure 2. *vip3-1* sepals emerge as larger primordia, but with WT-level variability in size (A) To estimate the size of abaxial primordia relative to the whole floral bud, a circle is fit on the bud, then the angle q formed by the center of the circle and the abaxial primordia ends (the sepal angle) is measured. (B) In relative to the whole floral buds, *vip3-1* abaxial primordia are wider than those in the WT. q is larger in the mutant (q_WT_ = 111° ± 2° and q_vip3-1_ = 144° ± 3°). However, the WT and the mutant have essentially the same coefficient of variation of q (∼ 8.5%). n = 20 sample for each genotype.

**Figure S3.**
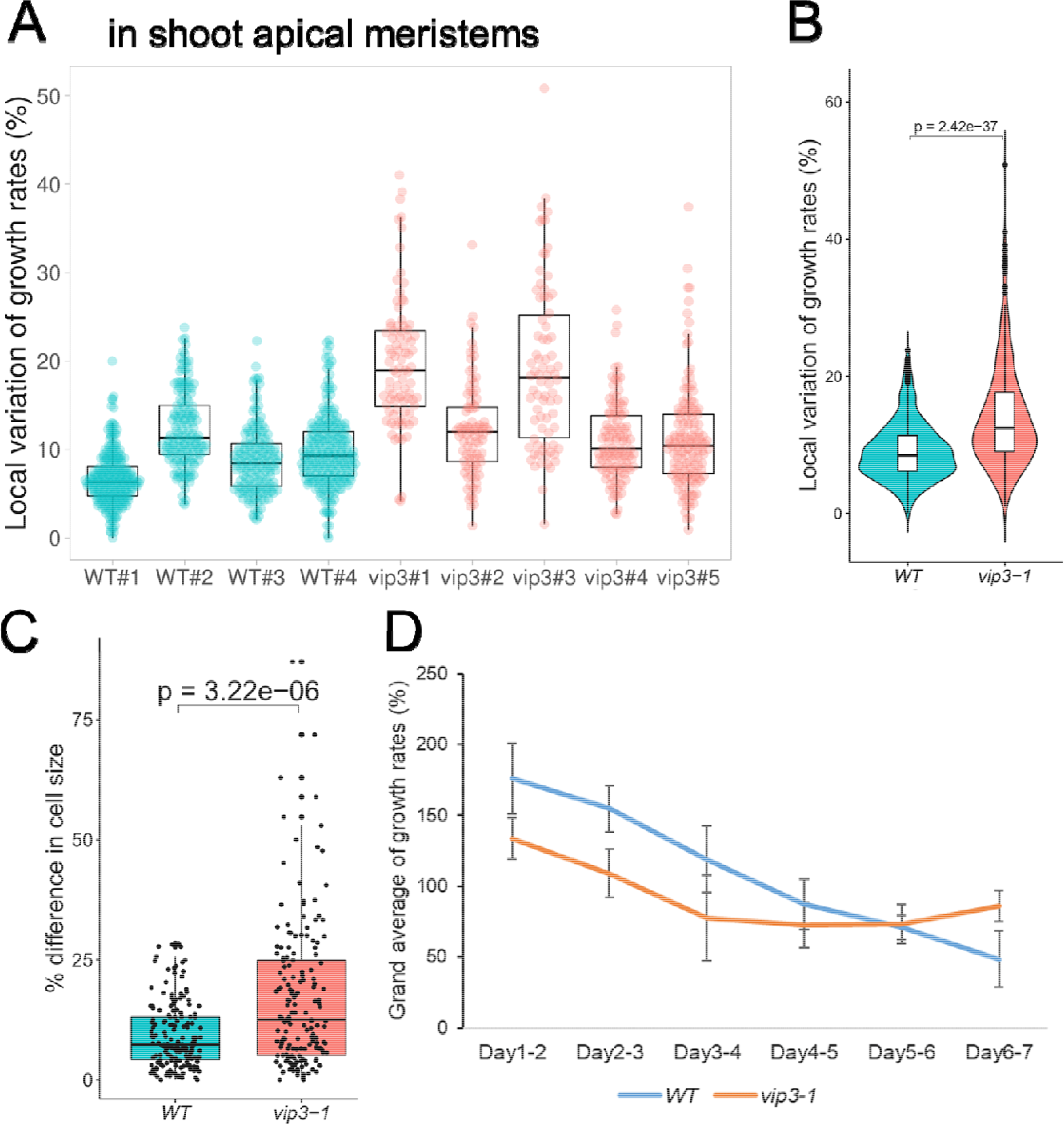
Related to Figure 3. Analysis of cell growth in *vip3-1* SAM, stomata and sepals (A) Local variation of growth rates in individual SAMs in WT and *vip3-1*. (B) Grand average of local variation of growth rates from individual samples in (A). (C) Difference (%) in cell size between two sister guard cells of the same stomata. n_WT_ = 171, n*_vip3-1_* = 169 pairs of guard cells, Kruskal–Wallis test. (D) Grand average of growth rates of WT and *vip3-1* sepals. Related to Figure 3G-H.

**Figure S4.**
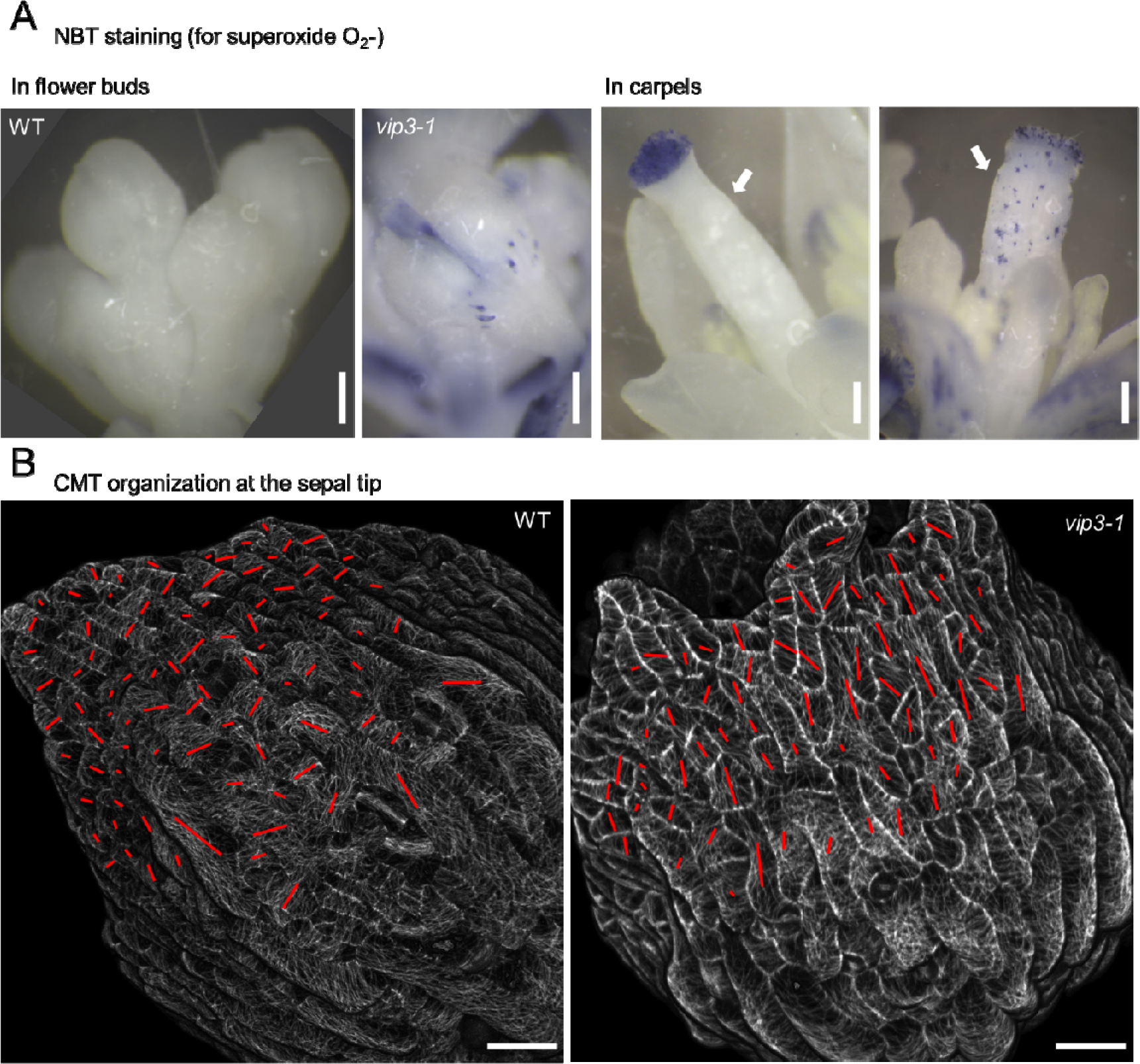
Related to Figure 4. Additional data on ROS and microtubule behavior in *vip3-1* (A) NBT staining for superoxide in WT (left panels) and *vip3-1* (right panels) inflorescences. Superoxide production in the mutant is higher in general and patchy in young flower buds as well as in the carpels (arrows). Scale bars: 0.1mm. (B) CMT organization at the sepal tip. Unlike the WT, most of tip cells in the mutant do not display a transverse pattern of CMTs. Scale bar: 20 µ m.

**Figure S5.**
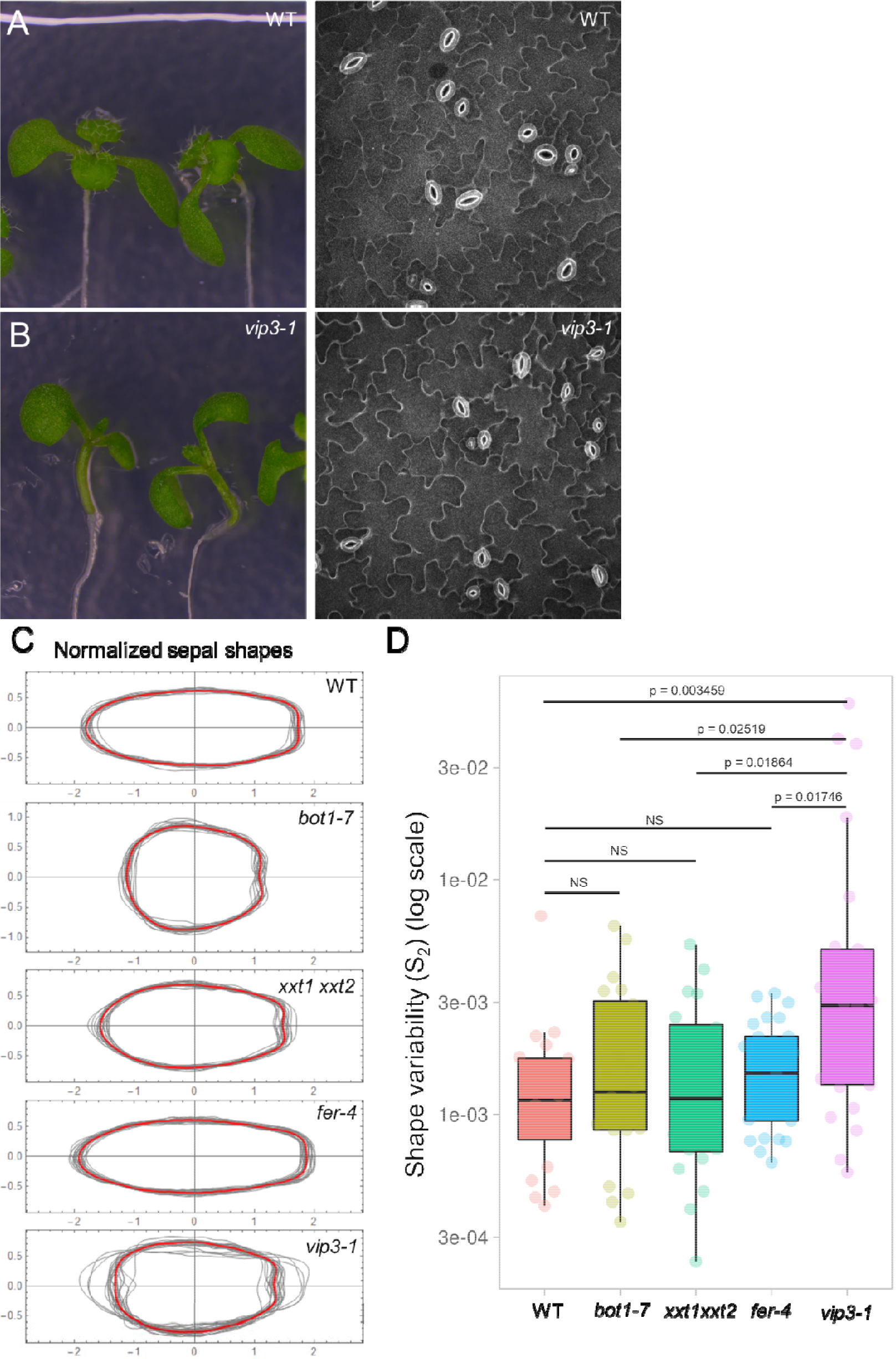
Related to Figure 5. (A,B) WT and *vip3-1* seedlings on normal agar (8g/L). (A) 9-day old WT seedlings on MS medium supplemented with 8g/L agar and their cells in the cotyledons under confocal microscopy (PI staining). (B) The same as (A) for *vip3-1*. Note that in this condition, no dead cells could be detected in the cotyledons. Scale bar = 1 mm (seedlings) and 100 µm (cells). (C,D) sepal shape variability in cell wall mutants. (C) Normalized contours of WT, *bot1-7, xxt1 xxt2, fer-4* and *vip3-1* abaxial sepals. The red outline is the average shape. (D) Quantification of shape variability (*S_2_*; log scale) with Kruskal–Wallis test. n = 20 for WT, *bot1-7* and *xxt1 xxt2*, = 25 for *fer-4* and *vip3-1*.

**Figure S6.**
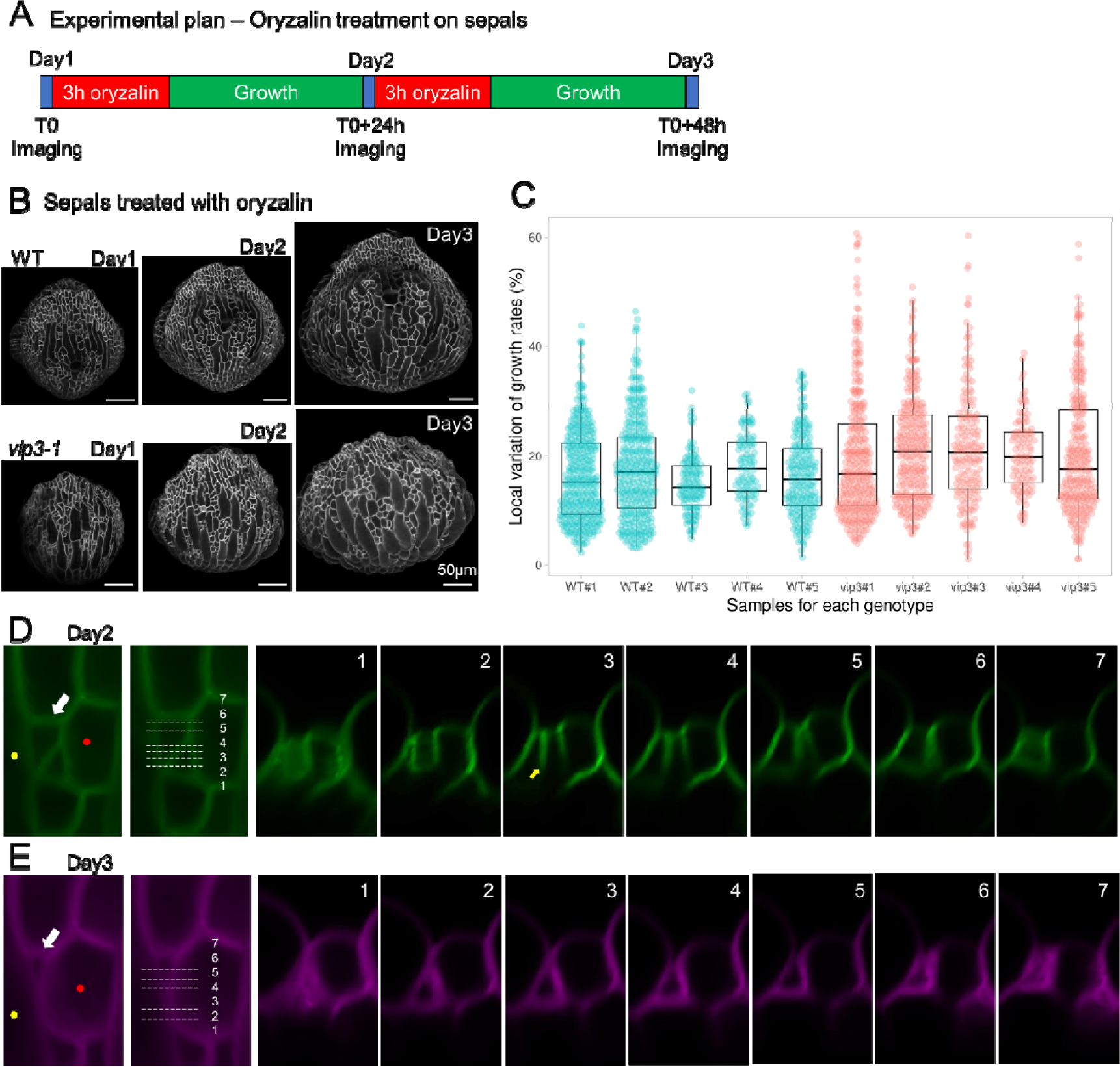
Related to Figure 6. Impacts of Oryzalin treatment of WT and *vip3* sepal growth (A) Experimental plan for the oryzalin treatment. Sepals were imaged and treated with oryzalin over 3 days as illustrated. (B) Representative images of WT and *vip3-1* sepals treated with Oryzalin 20 µg/L. Images were taken every 24 hours. Scale bars = 50 µ m. (C) Local variation of growth rates, calculated as coefficient of variation (CoV%), of cells in WT and *vip3-1* sepals treated with Oryzalin. Each boxplot represents a sample. (D-E) A series of orthogonal views of the squeezed cells (white arrow) in *vip3-1* along the white lines, at Day2 and Day3. The number on the orthogonal views corresponds to the white line on the left. At Day3, the volume of the squeezed cells is reduced. Note that the two cells indicated by the yellow and red dots were separated at Day2, but become almost contiguous at Day3.

## REFERENCES

Acar, M., Becskei, A., & van Oudenaarden, A. (2005). Enhancement of cellular memory by reducing stochastic transitions. Nature, 435(7039), 228–232. https://doi.org/10.1038/nature03524

Aegerter-Wilmsen, T., Aegerter, C. M., Hafen, E., & Basler, K. (2007). Model for the regulation of size in the wing imaginal disc of Drosophila. Mechanisms of Development, 124(4), 318–326. https://doi.org/10.1016/j.mod.2006.12.005

Aikawa, R., Nagai, T., Tanaka, M., Zou, Y., Ishihara, T., Takano, H., Hasegawa, H., Akazawa, H., Mizukami, M., Nagai, R., & Komuro, I. (2001). Reactive oxygen species in mechanical stress-induced cardiac hypertrophy. Biochemical and Biophysical Research Communications, 289(4), 901–907. https://doi.org/10.1006/bbrc.2001.6068

Ansel, J., Bottin, H., Rodriguez-Beltran, C., Damon, C., Nagarajan, M., Fehrmann, S., François, J., & Yvert, G. (2008). Cell-to-cell stochastic variation in gene expression is a complex genetic trait. PLoS Genetics, 4(4). https://doi.org/10.1371/journal.pgen.1000049

Antosz, W., Pfab, A., Ehrnsberger, H. F., Holzinger, P., Köllen, K., Mortensen, S. A., Bruckmann, A., Schubert, T., Längst, G., Griesenbeck, J., Schubert, V., Grasser, M., & Grasser, K. D. (2017). The composition of the arabidopsis RNA polymerase II transcript elongation complex reveals the interplay between elongation and mRNA processing factors. Plant Cell, 29(4), 854–870. https://doi.org/10.1105/tpc.16.00735

Araújo, I. S., Pietsch, J. M., Keizer, E. M., Greese, B., Balkunde, R., Fleck, C., & Hülskamp, M. (2017). Stochastic gene expression in Arabidopsis thaliana. Nature Communications, 8(1). https://doi.org/10.1038/s41467-017-02285-7

Bartschat, A., Hübner, E., Reischl, M., Mikut, R., & Stegmaier, J. (2016). XPIWIT—an XML pipeline wrapper for the Insight Toolkit. Bioinformatics, 32(2), 315–317. https://doi.org/10.1093/bioinformatics/btv559

Benikhlef, L., L’Haridon, F., Abou-Mansour, E., Serrano, M., Binda, M., Costa, A., Lehmann, S., & Métraux, J. P. (2013). Perception of soft mechanical stress in Arabidopsis leaves activates disease resistance. BMC Plant Biology, 13(1). https://doi.org/10.1186/1471-2229-13-133

Bichet, A., Desnos, T., Turner, S., Grandjean, O., & Höfte, H. (2001). BOTERO1 is required for normal orientation of cortical microtubules and anisotropic cell expansion in Arabidopsis. Plant Journal, 25(2), 137–148. https://doi.org/10.1046/j.1365-313X.2001.00946.x

Boudaoud, A., Burian, A., Borowska-Wykr t, D., Uyttewaal, M., Wrzalik, R., Kwiatkowska, D., & Hamant, O. (2014). FibrilTool, a^[^n ImageJ plug-in to quantify fibrillar structures in raw microscopy images. Nature Protocols, 9(2), 457–463. https://doi.org/10.1038/nprot.2014.024

Boulan, L., & Léopold, P. (2021). What determines organ size during development and regeneration? Development (Cambridge), 148(1), 0–3. https://doi.org/10.1242/dev.196063

Brás-Pereira, C., & Moreno, E. (2018). Mechanical cell competition. Current Opinion in Cell Biology, 51, 15–21. https://doi.org/10.1016/j.ceb.2017.10.003

Burk, D. H., Liu, B., Zhong, R., Morrison, W. H., & Ye, Z. H. (2001). A katanin-like protein regulates normal cell wall biosynthesis and cell elongation. Plant Cell, 13(4), 807–827. https://doi.org/10.1105/tpc.13.4.807

Burk, David H., & Ye, Z. H. (2002). Alteration of oriented deposition of cellulose microfibrils by mutation of a katanin-like microtubule-severing protein. Plant Cell, 14(9), 2145–2160. https://doi.org/10.1105/tpc.003947

Cavalier, D. M., Lerouxel, O., Neumetzler, L., Yamauchi, K., Reinecke, A., Freshour, G., Zabotina, O. A., Hahn, M. G., Burgert, I., Pauly, M., Raikhel, N. V., & Keegstra, K. (2008). Disrupting two Arabidopsis thaliana xylosyltransferase genes results in plants deficient in xyloglucan, a major primary cell wall component. Plant Cell, 20(6), 1519– 1537. https://doi.org/10.1105/tpc.108.059873

Chang, H. H., Hemberg, M., Barahona, M., Ingber, D. E., & Huang, S. (2008). Transcriptome-wide noise controls lineage choice in mammalian progenitor cells. Nature, 453(7194), 544–547. https://doi.org/10.1038/nature06965

Clough, S. J., & Bent, A. F. (1998). Floral dip: A simplified method for Agrobacterium-mediated transformation of Arabidopsis thaliana. Plant Journal, 16(6), 735–743. https://doi.org/10.1046/j.1365-313X.1998.00343.x

Crisucci, E. M., & Arndt, K. M. (2011). The Roles of the Paf1 Complex and Associated Histone Modifications in Regulating Gene Expression. Genetics Research International, 2011, 1–15. https://doi.org/10.4061/2011/707641

de Reuille, P. B., Routier-Kierzkowska, A. L., Kierzkowski, D., Bassel, G. W., Schüpbach, T., Tauriello, G., Bajpai, N., Strauss, S., Weber, A., Kiss, A., Burian, A., Hofhuis, H., Sapala, A., Lipowczan, M., Heimlicher, M. B., Robinson, S., Bayer, E. M., Basler, K., Koumoutsakos, P., … Smith, R. S. (2015). MorphoGraphX: A platform for quantifying morphogenesis in 4D. ELife, 4(MAY), 1–20. https://doi.org/10.7554/eLife.05864

Desai, R. V., Chen, X., Martin, B., Chaturvedi, S., Hwang, D. W., Li, W., Yu, C., Ding, S., Thomson, M., Singer, R. H., Coleman, R. A., Hansen, M. M. K., & Weinberger, L. S. (2021). A DNA repair pathway can regulate transcriptional noise to promote cell fate transitions. Science, 373(6557). https://doi.org/10.1126/science.abc6506

Dorcey, E., Rodriguez-Villalon, A., Salinas, P., Santuari, L., Pradervand, S., Harshman, K., & Hardtke, C. S. (2012). Context-dependent dual role of SKI8 homologs in mRNA synthesis and turnover. PLoS Genetics, 8(4), 1–9. https://doi.org/10.1371/journal.pgen.1002652

Dutilleul, C., Garmier, M., Noctor, G., Mathieu, C., Chétrit, P., Foyer, C. H., & de Paepe, R. (2003). Leaf Mitochondria Modulate Whole Cell Redox Homeostasis, Set Antioxidant Capacity, and Determine Stress Resistance through Altered Signaling and Diurnal Regulation. The Plant Cell, 15(5), 1212–1226. https://doi.org/10.1105/tpc.009464

Elowitz, M. B., Levine, A. J., Siggia, E. D., & Swain, P. S. (2002). Stochastic gene expression in a single cell. Science, 297(5584), 1183–1186. https://doi.org/10.1126/science.1070919

Fal, K., Cortes, M., Liu, M., Collaudin, S., Das, P., Hamant, O., & Trehin, C. (2019). Paf1c defects challenge the robustness of flower meristem termination in Arabidopsis thaliana . Development, dev.173377. https://doi.org/10.1242/dev.173377

Fal, K., Korsbo, N., Alonso-Serra, J., Teles, J., Liu, M., Refahi, Y., Chabouté, M.-E., Jönsson, H., & Hamant, O. (2021). Tissue folding at the organ–meristem boundary results in nuclear compression and chromatin compaction. Proceedings of the National Academy of Sciences, 118(8), e2017859118. https://doi.org/10.1073/pnas.2017859118

Fal, K., Liu, M., Duisembekova, A., Refahi, Y., Haswell, E. S., & Hamant, O. (2017). Phyllotactic regularity requires the Paf1 complex in Arabidopsis. Development, 144(23), 4428–4436. https://doi.org/10.1242/dev.154369

Feng, W., Kita, D., Peaucelle, A., Cartwright, H. N., Doan, V., Duan, Q., Liu, M. C., Maman, J., Steinhorst, L., Schmitz-Thom, I., Yvon, R., Kudla, J., Wu, H. M., Cheung, A. Y., & Dinneny, J. R. (2018). The FERONIA Receptor Kinase Maintains Cell-Wall Integrity during Salt Stress through Ca 2+ Signaling. Current Biology, 28(5), 666–675.e5. https://doi.org/10.1016/j.cub.2018.01.023

Fischl, H., Howe, F. S., Furger, A., & Mellor, J. (2017). Paf1 Has Distinct Roles in Transcription Elongation and Differential Transcript Fate. Molecular Cell, 65(4), 685–698.e8. https://doi.org/10.1016/j.molcel.2017.01.006

Guido, N. J., Lee, P., Wang, X., Elston, T. C., & Collins, J. J. (2007). A pathway and genetic factors contributing to elevated gene expression noise in stationary phase. Biophysical Journal, 93(11), L55–L57. https://doi.org/10.1529/biophysj.107.118687

Hamant, O., Das, P., & Burian, A. (2014). Time-Lapse Imaging of Developing Meristems Using Confocal Laser Scanning Microscope. In V. Žárský & F. Cvr ková (Eds.), Plant Cell Morphogenesis: Methods and Protocols (pp. 111–119). Humanča Press. https://doi.org/10.1007/978-1-62703-643-6_9

Hamant, O., Heisler, M. G., Jonsson, H., Krupinski, P., Uyttewaal, M., Bokov, P., Corson, F., Sahlin, P., Boudaoud, A., Meyerowitz, E. M., Couder, Y., & Traas, J. (2008). Developmental Patterning by Mechanical Signals in Arabidopsis. Science, 322(5908), 1650–1655. https://doi.org/10.1126/science.1165594

Heisler, M. G., Hamant, O., Krupinski, P., Uyttewaal, M., Ohno, C., Jönsson, H., Traas, J., & Meyerowitz, E. M. (2010). Alignment between PIN1 Polarity and Microtubule Orientation in the Shoot Apical Meristem Reveals a Tight Coupling between Morphogenesis and Auxin Transport. PLoS Biology, 8(10), e1000516. https://doi.org/10.1371/journal.pbio.1000516

Heitzler, P., & Simpson, P. (1991). The choice of cell fate in the epidermis of Drosophila. Cell, 64(6), 1083–1092. https://doi.org/10.1016/0092-8674(91)90263-X

Hervieux, N., Dumond, M., Sapala, A., Routier-Kierzkowska, A. L., Kierzkowski, D., Roeder, A. H. K., Smith, R. S., Boudaoud, A., & Hamant, O. (2016). A Mechanical Feedback Restricts Sepal Growth and Shape in Arabidopsis. Current Biology, 26(8), 1019–1028. https://doi.org/10.1016/j.cub.2016.03.004

Hong, L., Brown, J., Segerson, N. A., Rose, J. K. C., & Roeder, A. H. K. (2017). CUTIN SYNTHASE 2 Maintains Progressively Developing Cuticular Ridges in Arabidopsis Sepals. Molecular Plant, 10(4), 560–574. https://doi.org/10.1016/j.molp.2017.01.002

Hong, L., Dumond, M., Tsugawa, S., Sapala, A., Routier-Kierzkowska, A.-L., Zhou, Y., Chen, C., Kiss, A., Zhu, M., Hamant, O., Smith, R. S., Komatsuzaki, T., Li, C.-B., Boudaoud, A., & Roeder, A. H. K. (2016). Variable Cell Growth Yields Reproducible Organ Development through Spatiotemporal Averaging. Developmental Cell, 38(1), 15– 32. https://doi.org/10.1016/j.devcel.2016.06.016

Horiguchi, G., & Tsukaya, H. (2011). Organ size regulation in plants: Insights from compensation. Frontiers in Plant Science, 2(JUL), 1–6. https://doi.org/10.3389/fpls.2011.00024

Hou, L., Wang, Y., Liu, Y., Zhang, N., Shamovsky, I., Nudler, E., Tian, B., & Dynlacht, B. D. (2019). Paf1C regulates RNA polymerase II progression by modulating elongation rate. Proceedings of the National Academy of Sciences of the United States of America, 116(29), 14583–14592. https://doi.org/10.1073/pnas.1904324116

Hufnagel, L., Teleman, A. A., Rouault, H., Cohen, S. M., & Shraiman, B. I. (2007). On the mechanism of wing size determination in fly development. Proceedings of the National Academy of Sciences of the United States of America, 104(10), 3835–3840. https://doi.org/10.1073/pnas.0607134104

Jensen, G. S., Fal, K., Hamant, O., & Haswell, E. S. (2016). The RNA Polymerase-Associated Factor 1 Complex Is Required for Plant Touch Responses. Journal of Experimental Botany, 68(3), erw439. https://doi.org/10.1093/jxb/erw439

Kassambara, A. (2020a). Package ‘ggpubr.’ R Package Version 0.*1*, 6.

Kassambara, A. (2020b). rstatix: Pipe-friendly framework for basic statistical tests. R Package Version 0.*6*. 0.

Kierzkowski, D., Nakayama, N., Routier-Kierzkowska, A.-L., Weber, A., Bayer, E., Schorderet, M., Reinhardt, D., Kuhlemeier, C., & Smith, R. S. (2012). Elastic domains regulate growth and organogenesis in the plant shoot apical meristem. Science (New York, N.Y.), 335(6072), 1096–1099. https://doi.org/10.1126/science.1213100

Kimata, Y., Higaki, T., Kawashima, T., Kurihara, D., Sato, Y., Yamada, T., Hasezawa, S., Berger, F., Higashiyama, T., & Ueda, M. (2016). Cytoskeleton dynamics control the first asymmetric cell division in Arabidopsis zygote. Proceedings of the National Academy of Sciences of the United States of America, 113(49), 14157–14162. https://doi.org/10.1073/pnas.1613979113

Lampropoulos, A., Sutikovic, Z., Wenzl, C., Maegele, I., Lohmann, J. U., & Forner, J. (2013). GreenGate - A novel, versatile, and efficient cloning system for plant transgenesis. PLoS ONE, 8(12). https://doi.org/10.1371/journal.pone.0083043

Lampugnani, E. R., Khan, G. A., Somssich, M., & Persson, S. (2018). Building a plant cell wall at a glance. Journal of Cell Science, 131(2). https://doi.org/10.1242/jcs.207373

Landrein, B., Kiss, A., Sassi, M., Chauvet, A., Das, P., Cortizo, M., Laufs, P., Takeda, S., Aida, M., Traas, J., Vernoux, T., Boudaoud, A., & Hamant, O. (2015). Mechanical stress contributes to the expression of the STM homeobox gene in Arabidopsis shoot meristems. ELife, 4(DECEMBER2015), 1–27. https://doi.org/10.7554/eLife.07811

Legland, D., Arganda-Carreras, I., & Andrey, P. (2016). MorphoLibJ: Integrated library and plugins for mathematical morphology with ImageJ. Bioinformatics, 32(22), 3532–3534. https://doi.org/10.1093/bioinformatics/btw413

Little, S. C., Tikhonov, M., & Gregor, T. (2013). Precise Developmental Gene Expression Arises from Globally Stochastic Transcriptional Activity. Cell, 154(4), 789–800. https://doi.org/10.1016/j.cell.2013.07.025

Long, J. A., Moan, E. I., Medford, J. I., & Barton, M. K. (1996). A member of the KNOTTED class of homeodomain proteins encoded by the STM gene of Arabidopsis. Nature, 379(6560), 66–69. https://doi.org/10.1038/379066a0

Long, Y., Cheddadi, I., Mosca, G., Mirabet, V., Dumond, M., Kiss, A., Traas, J., Godin, C., & Boudaoud, A. (2020). Cellular Heterogeneity in Pressure and Growth Emerges from Tissue Topology and Geometry. Current Biology, 30(8), 1504–1516.e8. https://doi.org/10.1016/j.cub.2020.02.027

Malivert, A., Erguvan, Ö., Chevallier, A., Dehem, A., Friaud, R., Liu, M., Martin, M., Peyraud, T., Hamant, O., & Verger, S. (2021). FERONIA and microtubules independently contribute to mechanical integrity in the Arabidopsis shoot. PLOS Biology, 19(11), e3001454. https://doi.org/10.1371/journal.pbio.3001454

Meyer, H. M., Teles, J., Formosa-Jordan, P., Refahi, Y., San-Bento, R., Ingram, G., Jönsson, H., Locke, J. C. W., Roeder, A. H. K., Teles, J., Formosa-Jordan, P., Refahi, Y., Jönsson, H., Locke, J. C. W., San-Bento, R., & Ingram, G. (2017). Fluctuations of the transcription factor atml1 generate the pattern of giant cells in the arabidopsis sepal. ELife, 6, 1–41. https://doi.org/10.7554/eLife.19131

Milani, P., Gholamirad, M., Traas, J., Arnéodo, A., Boudaoud, A., Argoul, F., & Hamant, O. (2011). In vivo analysis of local wall stiffness at the shoot apical meristem in Arabidopsis using atomic force microscopy. Plant Journal, 67(6), 1116–1123. https://doi.org/10.1111/j.1365-313X.2011.04649.x

Nicholson, D. J. (2019). Is the cell really a machine? Journal of Theoretical Biology, 477, 108–126. https://doi.org/10.1016/j.jtbi.2019.06.002

Oh, S., Zhang, H., Ludwig, P., & Van Nocker, S. (2004). A mechanism related to the yeast transcriptional regulator Paf1c is required for expression of the Arabidopsis FLC/MAF MADS box gene family. Plant Cell, 16(11), 2940–2953. https://doi.org/10.1105/tpc.104.026062

Pan, Y., Heemskerk, I., Ibar, C., Shraiman, B. I., & Irvine, K. D. (2016). Differential growth triggers mechanical feedback that elevates Hippo signaling. Proceedings of the National Academy of Sciences of the United States of America, 113(45), E6974–E6983. https://doi.org/10.1073/pnas.1615012113

Park, Y. B., & Cosgrove, D. J. (2012). Changes in cell wall biomechanical properties in the xyloglucan-deficient xxt1/xxt2 mutant of Arabidopsis. Plant Physiology, 158(1), 465– 475. https://doi.org/10.1104/pp.111.189779

Postma, M., & Goedhart, J. (2019). Plotsofdata—a web app for visualizing data together with their summaries. PLoS Biology, 17(3), 1–8. https://doi.org/10.1371/journal.pbio.3000202

Preibisch, S., Saalfeld, S., & Tomancak, P. (2009). Globally optimal stitching of tiled 3D microscopic image acquisitions. Bioinformatics, 25(11), 1463–1465. https://doi.org/10.1093/bioinformatics/btp184

Raser, J. M., & O’Shea, E. K. (2004). Control of stochasticity in eukaryotic gene expression. Science, 304(5678), 1811–1814. https://doi.org/10.1126/science.1098641

Roeder, A. H. K. (2021). Arabidopsis sepals: A model system for the emergent process of morphogenesis. Quantitative Plant Biology, 2, e14. https://doi.org/10.1017/qpb.2021.12

Rué, P., Domedel-Puig, N., Garcia-Ojalvo, J., & Pons, A. J. (2012). Integration of cellular signals in chattering environments. Progress in Biophysics and Molecular Biology, 110(1), 106–112. https://doi.org/10.1016/j.pbiomolbio.2012.05.003

Rutowicz, K., Puzio, M., Halibart-Puzio, J., Lirski, M., Kotli ski, M., Krote, M. A., Knizewski, L., Lange, B., Muszewska, A., niegowska-ń wierk, K., Końcielniak, J., Iwanicka-Nowicka, R., Buza, K., JanowiakŚ, F., muda, ŚK., Jõesaar, I., śLaskowska-Kaszub, K., Fogtman, A., Kollist, H., … JerzmaŻnowski, A. (2015). A specialized histone H1 variant is required for adaptive responses to complex abiotic stress and related DNA methylation in Arabidopsis. Plant Physiology, 169(3), 2080–2101. https://doi.org/10.1104/pp.15.00493

Ryden, P., Sugimoto-Shirasu, K., Smith, A. C., Findlay, K., Reiter, W. D., & McCann, M. C. (2003). Tensile properties of Arabidopsis cell walls depend on both a xyloglucan cross- linked microfibrillar network and rhamnogalacturonan II-borate complexes. Plant Physiology, 132(2), 1033–1040. https://doi.org/10.1104/pp.103.021873

Sapala, A., & Smith, R. S. (2020). Osmotic Treatment for Quantifying Cell Wall Elasticity in the Sepal of Arabidopsis thaliana. In Plant Stem Cells: Methods and Protocols (Vol. 2094, pp. 101–112). https://doi.org/10.1007/978-1-0716-0183-9_11

Schürholz, A. K., López-Salmerón, V., Li, Z., Forner, J., Wenzl, C., Gaillochet, C., Augustin, S., Barro, A. V., Fuchs, M., Gebert, M., Lohmann, J. U., Greb, T., & Wolf, S. (2018). A comprehensive toolkit for inducible, cell type-specific gene expression in Arabidopsis. Plant Physiology, 178(1), 40–53. https://doi.org/10.1104/pp.18.00463

Shapiro, B. E., Tobin, C., Mjolsness, E., & Meyerowitz, E. M. (2015). Analysis of cell division patterns in the Arabidopsis shoot apical meristem. Proceedings of the National Academy of Sciences of the United States of America, 112(15), 4815–4820. https://doi.org/10.1073/pnas.1502588112

She, W., Grimanelli, D., Rutowicz, K., Whitehead, M. W. J., Puzio, M., Kotli ski, M., Jerzmanowski, A., & Baroux, C. (2013). Chromatin reprogramming duri^ń^ng the somatic-to-reproductive cell fate transition in plants. Development (Cambridge), 140(19), 4008– 4019. https://doi.org/10.1242/dev.095034

Shraiman, B. I. (2005). Mechanical feedback as a possible regulator of tissue growth. Proceedings of the National Academy of Sciences of the United States of America, 102(9), 3318–3323. https://doi.org/10.1073/pnas.0404782102

Smyth, D. R., Bowman, J. L., & Meyerowitz, E. M. (1990). Early flower development in Arabidopsis. Plant Cell, 2(8), 755–767. https://doi.org/10.1105/tpc.2.8.755

Stanislas, T., Hamant, O., & Traas, J. (2017). In-vivo analysis of morphogenesis in plants (Vol. 139, pp. 203–223). https://doi.org/10.1016/bs.mcb.2016.11.008

Strauss, S., Sapala, A., Kierzkowski, D., & Smith, R. S. (2019). Quantifying Plant Growth and Cell Proliferation with MorphoGraphX. In F. Cvr ková & V. Žárský (Eds.), Plant Cell Morphogenesis: Methods and Protocols (pp. 269–290). Springer New York. https://doi.org/10.1007/978-1-4939-9469-4_18

Team, R. C. (2013). R: A language and environment for statistical computing.

Tomson, B. N., & Arndt, K. M. (2013). The many roles of the conserved eukaryotic Paf1 complex in regulating transcription, histone modifications, and disease states. Biochimica et Biophysica Acta - Gene Regulatory Mechanisms, 1829(1), 116–126. https://doi.org/10.1016/j.bbagrm.2012.08.011

Torii, K. U. (2007). Stomatal patterning and guard cell differentiation. Plant Cell Monographs, 9(July), 343–359. https://doi.org/10.1007/7089_2007_135

Tsimring, L. S. (2014). Noise in biology. Reports on Progress in Physics, 77(2). https://doi.org/10.1088/0034-4885/77/2/026601

Tvergaard, V., & Needleman, A. (2018). Effect of Properties and Turgor Pressure on the Indentation Response of Plant Cells. *Journal of Applied Mechanics*, Transactions ASME, 85(6). https://doi.org/10.1115/1.4039574

Uyttewaal, M., Burian, A., Alim, K., Landrein, B., Borowska-Wykrt, D., Dedieu, A., Peaucelle, A., Ludynia, M., Traas, J., Boudaoud, A., Kwiatkowska, D., & Hamant, O. (2012). Mechanical stress acts via Katanin to amplify differences in growth rate between adjacent cells in Arabidopsis. Cell, 149(2), 439–451. https://doi.org/10.1016/j.cell.2012.02.048

Van Oss, S. B., Cucinotta, C. E., & Arndt, K. M. (2017). Emerging Insights into the Roles of the Paf1 Complex in Gene Regulation. Trends in Biochemical Sciences, 42(10), 788– 798. https://doi.org/10.1016/j.tibs.2017.08.003

Verger, S., Long, Y., Boudaoud, A., & Hamant, O. (2018). A tension-adhesion feedback loop in plant epidermis. ELife, 7, 1–25. https://doi.org/10.7554/eLife.34460

Weinberger, L. S., Burnett, J. C., Toettcher, J. E., Arkin, A. P., & Schaffer, D. V. (2005). Stochastic gene expression in a lentiviral positive-feedback loop: HIV-1 Tat fluctuations drive phenotypic diversity. Cell, 122(2), 169–182. https://doi.org/10.1016/j.cell.2005.06.006

Yanofsky, M. F., Ma, H., Bowman, J. L., Drews, G. N., Feldmann, K. A., & Meyerowitz, E. M. (1990). The protein encoded by the Arabidopsis homeotic gene agamous resembles transcription factors. Nature, 346(6279), 35–39. https://doi.org/10.1038/346035a0

Zhang, H., Ransom, C., Ludwig, P., & Van Nocker, S. (2003). Genetic analysis of early flowering mutants in Arabidopsis defines a class of pleiotropic developmental regulator required for expression of the flowering-time switch FLOWERING LOCUS C. Genetics, 164(1), 347–358.

Zhao, F., Chen, W., Sechet, J., Martin, M., Bovio, S., Lionnet, C., Long, Y., Battu, V., Mouille, G., Monéger, F., & Traas, J. (2019). Xyloglucans and Microtubules Synergistically Maintain Meristem Geometry and Phyllotaxis. Plant Physiology, 181(3), 1191–1206. https://doi.org/10.1104/pp.19.00608

Zhu, M., Chen, W., Mirabet, V., Hong, L., Bovio, S., Strauss, S., Schwarz, E. M., Tsugawa, S., Wang, Z., Smith, R. S., Li, C. B., Hamant, O., Boudaoud, A., & Roeder, A. H. K. (2020). Robust organ size requires robust timing of initiation orchestrated by focused auxin and cytokinin signalling. Nature Plants. https://doi.org/10.1038/s41477-020-0666-7

